# A *gαq* signaling pathway in dopaminergic neurons suppresses memory

**DOI:** 10.64898/2026.10.13.758534

**Authors:** Juan C. Martinez-Cervantes, Prachi Shah, Isaac Cervantes-Sandoval

## Abstract

Which experiences persist as memories is a fundamental question in neurobiology. The balance between memory persistence and suppression is thought to be optimal for an organism to adapt to the ever-changing environment. Memory suppression is a process by which the brain limits the strength and expression of memories. Memory suppressors are genes whose disruption can enhance memory at different stages: acquisition, consolidation, forgetting, or recall. Here, using aversive classical conditioning in *Drosophila,* we show that Gαq acts as a memory suppressor in PPL1 dopaminergic neurons. Knockdown of *gαq* in dopaminergic neurons enhances acquisition, strengthens anesthesia-resistant memory, and impairs reversal learning. In vivo functional imaging revealed that *gαq* knockdown amplifies shock-evoked responses in a type of PPL1 neuron. Additionally, using *ex vivo* imaging and pharmacological experiments, we reveal that both Gαq and the OAMB receptor alter octopamine-evoked ER calcium responses in PPL1 neurons. Our results uncover a new role of Gαq signaling within dopaminergic neurons that suppresses memory.

## Introduction

Determining which memories, and which aspects of those memories, are stored in the brain is crucial for predicting future outcomes. Memory suppression is the process by which the brain limits the strength and expression of memories. Memory suppressors are genes, and their encoded proteins, whose reduced expression enhances memory at distinct stages of memory processing, including acquisition, consolidation, forgetting, and recall (Noyes et al., 2021).

Experimental evidence indicates that some memory suppressors promote behavioral flexibility in dynamic environments by attenuating outdated or maladaptive associations (Hoeffer et al., 2008; Shuai et al., 2010, 2011; Berry et al., 2012; Cervantes-Sandoval et al., 2016; Dong et al., 2016; Zhang et al., 2016, 2018; Davis and Zhong, 2017; Gao et al., 2019; Mariottini et al., 2019; Phan et al., 2019; Noyes et al., 2021). For example, disrupting forgetting impairs reversal-learning and extinction performance, indicating reduced behavioral flexibility. Strikingly, inhibiting other memory suppressors, such as Hdac2, has been reported to enhance both memory and flexibility in reversal and extinction assays (Guan et al., 2009). These seemingly opposing effects remain poorly understood, though one proposed explanation is that suppressors acting early in memory processing—such as during acquisition—may promote flexibility through a mechanism distinct from those acting during forgetting. Resolving these opposing effects will likely require circuit-specific dissection, underscoring that an appropriate balance of memory suppression is essential for optimizing adaptive behavior.

In *Drosophila*, the mushroom body, and in particular, the dopaminergic neuronal population innervating this structure, constitutes a well-established circuit regulating both the encoding and forgetting of aversive memories. PPL1 dopaminergic neurons (DANs) convey punishment signals through reinforcement-driven evoked responses during learning (Schwaerzel et al., 2003; Claridge-Chang et al., 2009; Aso et al., 2010, 2012, 2020; Burke et al., 2012; Liu et al., 2012; Aso and Rubin, 2016). Conversely, dopamine signaling that occurs after learning, and is modulated by the behavioral state of the animal, can reduce memory expression (Berry et al., 2015, 2018; Cohn et al., 2015; Handler et al., 2019; Sabandal et al., 2021). Molecular evidence suggests that these divergent processes are mediated by two distinct dopamine receptors (Kim et al., 2007; Berry et al., 2012; Himmelreich et al., 2017; Handler et al., 2019) located postsynaptically in the Kenyon cells (KCs) of the mushroom body (Perez-Orive et al., 2002; Honegger et al., 2011). The D1-like receptor Dop1R1 mediates memory encoding by sensing dopamine levels during reinforcement, whereas dopamine signals convey memory suppression through the Dop1R2 (DAMB) receptor. Downstream of DAMB, several proteins in KCs have been implicated in the same memory-suppression pathway, including Rac1, Scribble, and Gαq (Shuai et al., 2010; Cervantes-Sandoval et al., 2016; Himmelreich et al., 2017). DAMB preferentially couples to Gαq to trigger Ca²⁺ signaling, and knocking down *gαq* in KCs decreases forgetting with no obvious effect on acquisition (Himmelreich et al., 2017).

In this study, we show that Gαq also functions as a memory suppressor within a subset of PPL1 dopaminergic neurons, regulating acquisition, forgetting, memory flexibility, and anesthesia-resistant memory. We further demonstrate that multiple G-protein-coupled receptors (GPCRs) modulate memory suppression in PPL1 neurons, and we link these behavioral effects to dopaminergic functional imaging.

## Results and discussion

Synaptic scaffolding molecules, which by virtue of their ability to simultaneously bind several proteins, play crucial roles in arranging the structural and functional complexes for signaling. Scribble, one of these proteins, has been implicated in memory suppression of olfactory memory in *Drosophila* (Cervantes-Sandoval et al., 2016). We hypothesized that Scribble organizes a signaling complex required for memory suppression. To identify Scribble-binding partners, we used the endogenous protein trap *scrb::GFP^CA07683^* (Buszczak et al., 2007) as bait for anti-GFP immunoprecipitation (IP) from adult head lysates followed by mass spectrometry (Fig. 1A and S1).

**Figure 1.**
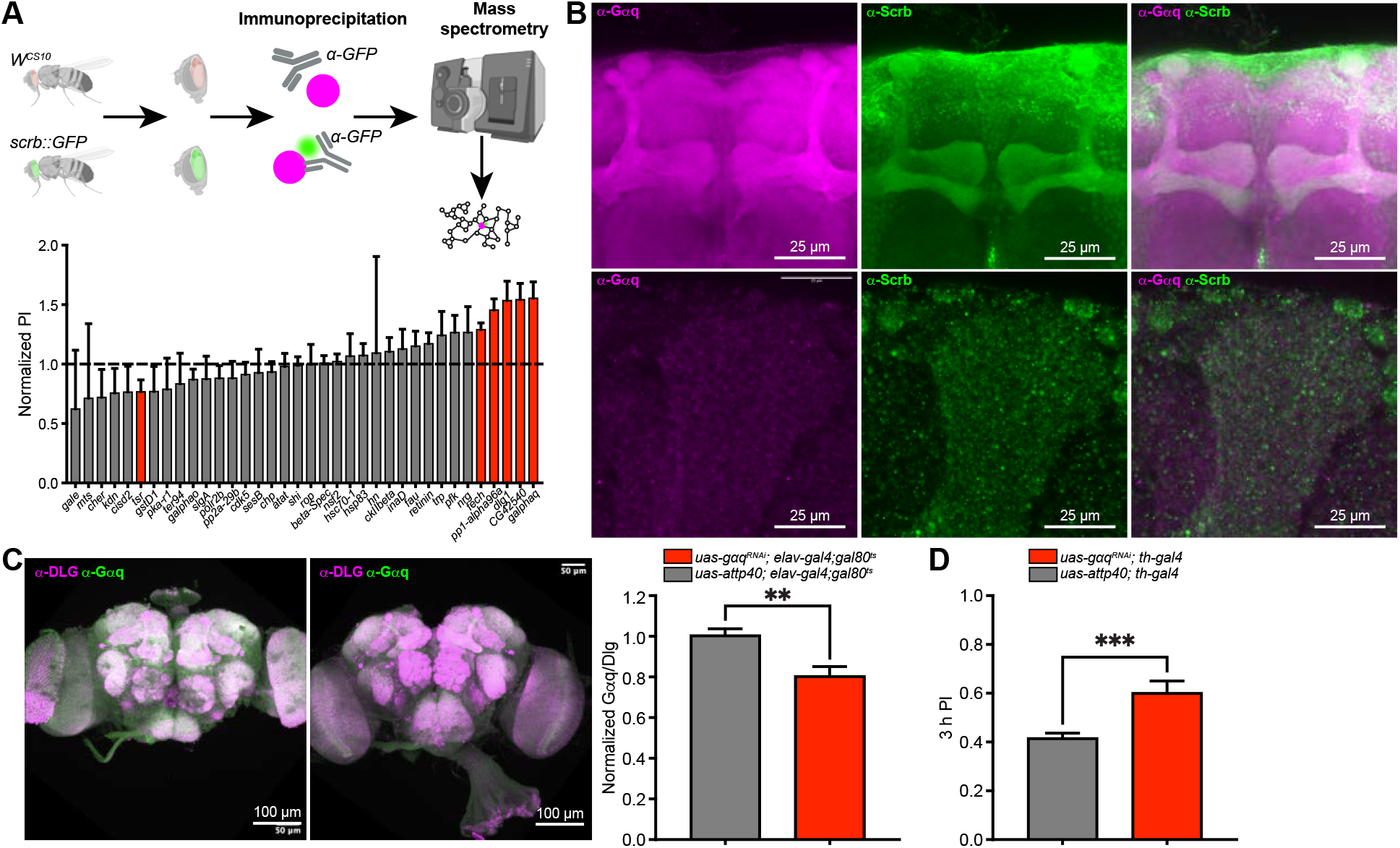
Identification of candidate memory-suppression genes using a behavioral screen. (A) Normalized 3 h memory performance scores of candidates after aversive olfactory conditioning. Each experimental group was normalized to its genetic control tested in parallel. Mann-Whitney test comparing the knockdown against its genetic control. Red bars indicate significance, *p < 0.05. n ≥ 6*. Data are shown as mean ± SEM. (B) Top. Maximum-intensity projections of confocal stacks showing endogenous Gαq (left; anti-Gαq) and Scribble (anti-Scribble) in the mushroom bodies; merge (right). Bottom: Maximum-intensity projection of an expansion microscopy stack in the mushroom body lobe, α3 compartment. Gαq and Scribble signal in α3 neuropil shows partial colocalization. (C) Top. Representative images of brains showing expression of Gαq and Dlg in *gαq^RNAi^* and genetic controls using the *elav-gal4; gal80^ts^* driver after RNAi induction during adulthood. Bottom. Normalized expression of Gαq relative to Dlg in *gαq^RNAi^* fly brains. *\*p = 0.0059. n = 7–8*. Data are shown as mean ± SEM. (D) Knocking down *gαq* in dopaminergic neurons using the *th-gal4* enhanced 3 h aversive memory. Mann-Whitney test, *\*p = 0.0006. n = 11*. Data are shown as mean ± SEM.

Across four biological replicates, Scribble was the most abundant protein recovered (2845–3516 spectral counts), indicating robust enrichment of the bait. Among the filtered candidates, we detected the canonical polarity partner Dlg1 (3/4 replicates) and multiple signaling and cytoskeletal proteins, including Gαq (4/4 replicates) and Cofilin (3/4 replicates), consistent with Scribble coupling to actin-regulatory pathways previously implicated in memory suppression. Additional recurrent proteins included phosphatase/kinase-associated factors and vesicle/trafficking components, alongside housekeeping proteins expected from head-derived preparations. The co-purification of InaD alongside InaC and Trp, which together constitute the InaD-organized phototransduction complex (Tsunoda et al., 1997), demonstrates that the IP captures intact multiprotein assemblies, providing internal validation (Fig. S1).

Proteomic identification of Scribble-associated proteins gave us a set of candidate members of a memory-suppression pathway. We tested the 36 most enriched hits and conducted a candidate-based RNAi screen using pan-neuronal expression of each RNAi line (*uas-dcr2; nsyb-gal4*) measuring aversive olfactory memory performance 3 h after conditioning (Fig. 1A). We used a pan-neuronal driver because Scribble has been implicated in memory suppression in both Kenyon cells (KC) of the mushroom body (MB) and PPL1 dopaminergic neurons that innervate the MB. This approach allowed us to identify candidate genes whose knockdown affects memory through either neuronal population without presupposing the cellular locus of the Scribble-associated pathway. In this screen, RNAi knockdown of *fech*, *pp1-α96a*, *dlg1*, *cg42540*, and *gαq* significantly increased 3 h aversive olfactory memory, whereas knockdown of twinstar (cofilin/tsr) reduced memory. Among all hits, *gαq* produced the strongest enhancement.

To visualize Gαq in the adult brain, we performed immunohistochemistry using an anti-Gαq antibody. Gαq immunoreactivity was detected in multiple brain regions, including the MB, where signal was enriched in all three lobes, α/β, α’/β’ and γ (Fig. 1B). We then performed expansion microscopy (ExM) to improve resolution. ExM of Gαq revealed a punctate signal across the brain and especially enriched in the MB lobes. Expansion microscopy also revealed that Gαq and Scribble signals overlap within the MB lobes (Fig. 1B, bottom).

Using the same antibodies, we validated the efficiency of the *gαq* RNAi transgene line by quantifying the ratio of anti-Gαq signal relative to anti-Dlg1. Pan-neuronal *gαq* knockdown only during adulthood using the TARGET system showed significantly reduced Gαq signal compared to controls (Fig. 1C).

Multiple memory suppressor genes found in *Drosophila* have been shown to function in both KCs and DANs, including Scribble, Stromalin, Sickie and others (Cervantes-Sandoval et al., 2016; Phan et al., 2019; Zhang et al., 2022). Because Gαq has a previously established role in KC for memory suppression (Himmelreich et al., 2017), we tested its function in PPL1-DANs. Knockdown of *gαq* using *th-gal4* enhanced memory performance 3 h post-training (Fig. 1D). Odor avoidance and shock reactivity were normal in *gαq^RNAi^*flies (Fig. S2).

## Gαq regulates acquisition and behavioral flexibility

To confirm the memory enhancement effect, we used two independent RNAi transgenes to knock down *gαq* in DANs. Using transgenes on the second and third chromosomes that target *gαq*, we recapitulated the 3 h post-training enhanced memory effect (Fig. 2A). Then, we analyzed the full memory retention curve to try to pinpoint when the effect of enhanced memory becomes apparent. We observed that *gαq^RNAi^* enhances the behavioral expression of aversive olfactory memory from the immediate memory (3 min memory) up to 24 h memory (Fig. 2B). These data suggest that *gαq* knockdown in DANs increases memory acquisition since immediate memory is significantly enhanced. To test this directly and eliminate the possibility of ceiling-effect confounds we performed two different acquisition experiments by training the flies with an increasing number of shock pulses or 12 pulses of increasing shock intensity. Results showed a memory enhancement in *gαq^RNAi^*flies compared to controls when they received 6 or 12 shocks per training (Fig. 2C). In addition, no significant effects were observed when changing the voltage during conditioning except for 90 V (Fig. 2D). These results suggest that the memory suppression effect caused by KD of *gαq* might result from enhanced learning.

**Figure 2.**
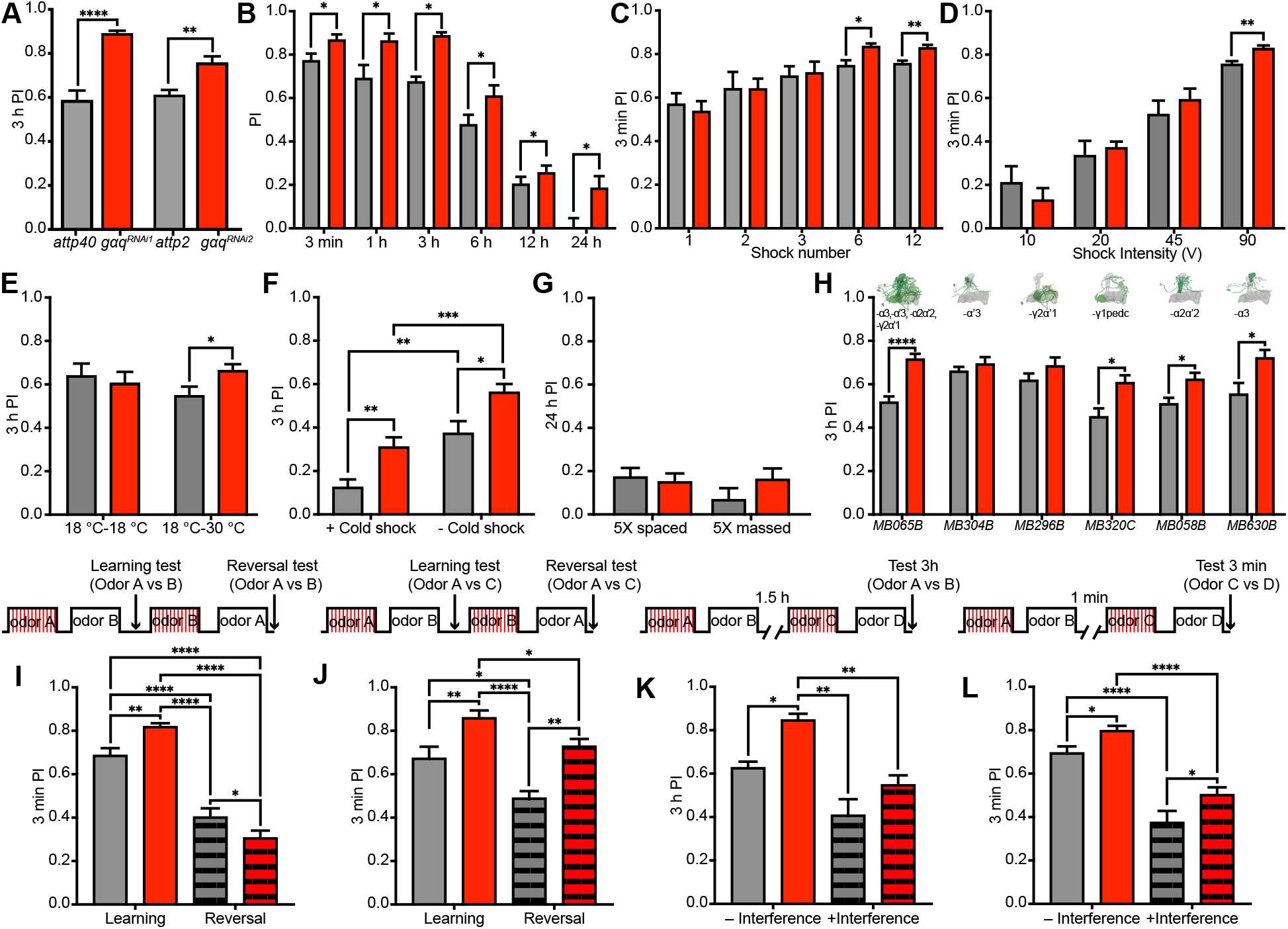
Knocking down *gαq* in DANs enhances memory performance. (A) Knocking down *gαq* in DANs using *MB504B* and *th-gal4* with two independent RNAi transgenes enhances 3 h memory. Mann-Whitney test. RNAi line 1: *\*\*\*p = 0.0006, n = 11*. RNAi line 2: *\*\*p = 0.0036, n = 8-14*. Data are shown as mean ± SEM. (B) Memory performance of *gαq^RNAi^* in the *th-gal4* driver is enhanced during acquisition and up to 24 h after aversive conditioning. Statistics: two-tailed Mann-Whitney tests comparing genotypes at each time point. *\*p < 0.05. n = 6-15*. Data are shown as mean ± SEM. (C) Immediate memory is enhanced in *gαq^RNAi^* in training protocols using 6 or 12 electric shock pulses. Mann-Whitney tests, 6 shocks: *\*\*p = 0.0127, n = 12-14*. 12 shocks: *\*\*p = 0.0025, n = 11-13*. Data are shown as mean ± SEM. (D) Varying the shock voltage in the training protocols does not affect immediate memory performance except for 90 V. Mann-Whitney tests. *\*\*p = 0.0025, n = 6-13*. Data are shown as mean ± SEM. (E) Knocking down *gαq* only during adulthood using the TARGET system in *th-gal4* cells enhances 3 h memory performance. Mann-Whitney test. *\*p = 0.032. n = 15-17*. Data are shown as mean ± SEM. (F) Knocking down *gαq* in the dopaminergic neurons increases both anesthesia-sensitive and anesthesia-resistant memory. Two-way ANOVA followed by Fisher’s LSD post hoc comparisons: + Cold shock (control vs *gαq^RNAi^*), *\*\*p = 0.0025*; –Cold shock (control vs *gαq^RNAi^*) *p = 0.034. n = 6-16*. Data are shown as mean ± SEM. (G) Knocking down *gαq* in the dopaminergic neurons does not alter protein synthesis-dependent LTM or long-term anesthesia resistant memory. Mann-Whitney tests. *n = 10*. Data are shown as mean ± SEM. (H) Knocking down *gαq* in subsets of PPL1 neurons recapitulates the memory enhancement 3 h after conditioning. Statistics: two-tailed Mann-Whitney tests comparing genotypes at each time point. Data are shown as mean ± SEM. (I) Memory performance in MB504B>*gαq^RNAi^*is higher than control after learning. In contrast, memory performance is significantly lower after reversal learning to the updated contingency. Two-way ANOVA, *\*\*\*\*p < 0.0001*; Holm–Šídák post hoc comparison of reversal performance (control vs MB504B> *gαq^RNAi^*), *\*p = 0.0180; n = 7-8*. Data are shown as mean ± SEM. (J) Memory performance in MB504B> *gαq^RNAi^* is higher after both learning and reversal learning in a paradigm in which a third novel odor is introduced during test. Two-way ANOVA, Holm–Šídák post hoc comparison test. \*\**p = 0.0026. n = 6-8*. Data are shown as mean ± SEM. (K) MB504B>*gαq^RNAi^*flies show similar memory decay after retroactive interference compared to controls. Two-way ANOVA across conditions, \*\*\*\**p = 0.0007*. Data are shown as mean ± SEM. (L) Proactive interference decreased memory performance in both control and MB504B> *gαq^RNAi^*flies, but knockdown flies retained higher memory scores than controls under interference. Two-way ANOVA followed by Holm–Šídák multiple comparisons for proactive interference groups adjusted *\*p = 0.0185; n = 7-8*. Data are shown as mean ± SEM.

Gαq is a major intracellular signaling pathway downstream of GPCR activation, and numerous GPCRs are thought to signal through Gαq. Thus, constitutive knockdown of *gαq* during development could produce broad developmental effects that might confound the interpretation of the observed memory phenotype. To determine whether the memory-enhancement phenotype results from developmental perturbations, we used the TARGET system to restrict *gαq^RNAi^* expression to adulthood. Adult-specific knockdown of *gαq* was sufficient to enhance memory retention 3 h after training, indicating that the phenotype does not require developmental disruption. In contrast, no significant differences in memory were observed in flies maintained at the non-inducing temperature, which prevents *gαq^RNAi^* expression (Fig. 2E).

In *Drosophila*, aversive olfactory conditioning produces at least two distinguishable, parallel forms of memory (Quinn and Dudai, 1976; Tully et al., 1994). Anesthesia-sensitive memory (ASM) is a short-lived, labile memory component, whereas anesthesia-resistant memory (ARM) might persist for at least 24 h and is resistant to several forms of anesthesia (Folkers et al., 1993; Shah et al., 2026). To determine which memory component is affected by *gαq* knockdown, we trained flies and subjected them to cold-shock anesthesia 2 h after training, followed by a memory test 3 h post-conditioning. As expected, *gαq^RNAi^*flies exhibited enhanced memory compared with genetic controls in the absence of cold-shock anesthesia. Notably, *gαq^RNAi^* flies that received cold shock 2 h after training also exhibited enhanced memory compared with genetic controls, indicating that at least a portion of the memory enhancement produced by *gαq* knockdown is resistant to cold-shock anesthesia. However, memory performance remained significantly higher in *gαq^RNAi^*flies that did not receive cold shock compared with those that did, suggesting that another component of the enhanced memory is sensitive to cold-shock anesthesia. Together, these results suggest that *gαq* knockdown enhances both anesthesia-resistant and anesthesia-sensitive memory components (Fig. 2F). Interestingly, when flies were trained using a 5× massed training protocol and memory was assessed 24 h later, no significant effect of *gαq* knockdown was detected. Similarly, the 5× spaced training protocol, which typically promotes protein synthesis-dependent long-term memory, did not reveal a significant difference between genotypes (Fig. 2G). Together, these results suggest that *gαq* knockdown does not enhance the consolidation of long-term memory under these training conditions.

Interference-based memory suppression refers to forgetting that results from competing information acquired before or after a relevant experience (Shuai et al., 2010; Cervantes-Sandoval et al., 2016; Zhao et al., 2023). We performed four experiments to determine whether *gαq* knockdown affects forgetting induced by different forms of memory interference. First, we tested the ability of flies to update a previously learned memory by using a reversal-learning paradigm. Flies were first trained with one odor contingency (CS+ = A, CS− = B) and immediately retrained with the opposite contingency (CS+ = B, CS− = A). Memory was then assessed using the same odors presented during training (Fig. 2I). Control flies showed a substantial avoidance to the odor most recently paired with the shock during the reversal training, consistent with their ability to update the original odor association by suppressing or weakening the previously learned contingency. In contrast, *gαq^RNAi^* flies showed significantly impaired performance for the second contingency compared with genetic controls (Fig. 2I), suggesting that *gαq* knockdown impairs the ability to fully update previously established memory representations.

We next directly assessed the persistence of the original memory following the competing training experience. Flies were trained using the same reversal-learning protocol but were subsequently tested with the original CS+ paired against a novel, never-experienced odor (A vs. C). In control flies, the second training session produced significant forgetting of the original memory, consistent with interference from the competing contingency. However, *gαq^RNAi^*flies retained significantly stronger memory for the original contingency than genetic controls (Fig. 2J). Despite this, when the initial memory in *gαq^RNAi^* flies is compared within genotype, there is a significant reduction of memory performance after reversal learning. These data indicate that reversal learning is not fully impaired. The last two experiments together, suggest that *gαq* knockdown only partially impairs forgetting induced by competing information and consequently only partially reduces the ability to update or suppress previously established memories.

We then examined retroactive interference (Shuai et al., 2010; Cervantes-Sandoval et al., 2016), in which a subsequent learning experience reduces memory for a previously acquired association. In this paradigm, flies received a second training session 1.5 h after the initial training using a completely different odor combination (odors A/B vs C/D), and memory for the original association was assessed 3 h later. *gαq^RNAi^* flies still exhibited a trend toward enhanced memory performance when compared to the genetic control (p = 0.084). Nevertheless, *gαq* knockdown flies, exhibited a significant reduction in memory following the second training session, indicating that retroactive interference remained intact (Fig. 2K). Thus, *gαq* knockdown does not appear to impair forgetting produced by an unrelated subsequent memory.

Finally, we tested proactive interference, in which a prior learning experience interferes with the acquisition or expression of a subsequent memory. Flies were first trained with an A/B odor combination and subsequently trained with a distinct C/D combination, after which memory for the second contingency was assessed (C/D). Control flies exhibited significantly reduced memory following the prior training experience compared with flies that did not receive the initial training, demonstrating robust proactive interference (Fig. 2L). *gαq^RNAi^* flies similarly exhibited proactive-interference-induced memory suppression (Fig. 2L). These results indicate that the genotype difference is unlikely to reflect a general resistance to interference, but rather an enhancement in baseline memory acquisition that persists in the presence of proactive interference.

Together, these findings indicate that *gαq* knockdown produces a selective and mild alteration in interference-dependent memory suppression. The enhanced retention of the original memory and impaired reversal learning are consistent with a reduced ability to suppress or update previously established memories, whereas retroactive and proactive interference remain largely intact. Thus, *gαq* knockdown does not produce a general impairment in forgetting but instead appears to selectively affect behavioral flexibility and the suppression of fully competing memory representations (Fig. 2). Combined with the enhanced anesthesia-resistant memory described above, these results suggest that Gαq signaling in dopaminergic neurons contributes to the regulation of memory strength, retention, and specific forms of memory suppression.

## Gαq behaviorally maps out to a subset of Dopaminergic neurons

The mushroom body is organized into anatomically and functionally distinct compartments that serve as discrete units of associative plasticity (Aso et al., 2014; Aso and Rubin, 2016). Each compartment is defined by the convergence of KC axons and the dendrites of specific MB output neurons (MBONs), with synaptic strength locally modulated by stereotyped dopaminergic inputs. Learning-induced plasticity at KC>MBON synapses is therefore expressed in a compartment-specific manner, allowing individual compartments to encode distinct components of memory with different dynamics and stability (Aso and Rubin, 2016). This modular organization enables parallel, and sometimes opposing, influences on behavior to be integrated at the level of MB output.

To determine which PPL1-DAN compartments require Gαq signaling to regulate memory, we selectively knocked down *gαq* in individual PPL1 dopaminergic neuron populations using specific split-*gal4* drivers. We first used a PPL1-DAN driver that selectively targets the four PPL1-DANs that innervate the MB lobes: PPL1-α3, PPL1-α′3, PPL1-α′2α2, and PPL1-γ2α′1, without expression in other DAN populations. Knockdown of *gαq* with this driver phenocopied the enhanced-memory phenotype observed with the broader *th-gal4* driver (Fig. 2H), which is expressed throughout the PPL1 population as well as in additional dopaminergic clusters, including DANs outside the MB. We next restricted *gαq* knockdown to individual PPL1 compartments to determine which neurons contribute to this phenotype. Knockdown in PPL1-α′3 or PPL1-γ2α′1 did not significantly affect memory performance (Fig. 2H). In contrast, knockdown in PPL1-γ1pedc, PPL1-α′2α2, or PPL1-α3 significantly enhanced memory performance (Fig. 2H). These results identify a subset of PPL1-DAN compartments in which Gαq signaling contributes to the suppression of aversive memory.

## Gαq modulates reinforcement signals during conditioning

During aversive conditioning, PPL1 dopaminergic neurons provide reinforcement signals to defined compartments of the MB vertical lobes (Claridge-Chang et al., 2009; Mao and Davis, 2009; Aso et al., 2010; Burke and Waddell, 2011; Huang et al., 2024). Activation of these neurons instructs synaptic plasticity at KC>MBON synapses and contributes to the formation of aversive memory (Séjourné et al., 2011; Hige et al., 2015; Owald et al., 2015; Berry et al., 2018; Cervantes-Sandoval et al., 2020). In addition to their role in memory formation, subsets of PPL1 neurons contribute to memory suppression and updating (Berry et al., 2012, 2015; Sabandal et al., 2021). PPL1 DANs also respond to odors innately and encode learned valence in a heterogeneous and bidirectional manner following conditioning (Cervantes-Sandoval et al., 2017; Siju et al., 2020; Huang et al., 2024). Thus, compartment-specific dopaminergic signaling by PPL1 neurons plays a central role in the encoding, maintenance, and modification of aversive memories over time (Aso et al., 2014; Aso and Rubin, 2016).

Because Gαq signaling can promote intracellular Ca²⁺ mobilization, we hypothesized that reducing *gαq* expression in PPL1 DANs would alter Ca²⁺ dynamics at their synaptic terminals, where reinforcement signals are delivered to the MB. To test this possibility, we performed in vivo functional calcium imaging by co-expressing GCaMP6f and *gαq^RNAi^* in PPL1 DANs and recording Ca²⁺ activity from PPL1-α3 terminals in head-fixed, awake flies (Fig. 3A–B). We first measured olfactory responses to 4-MCH and 3-OCT and found no significant differences between control and *gαq^RNAi^*flies for either odor (Fig. 3C). Odor-evoked responses in PPL1-α3 were relatively small compared with those observed in other PPL1 neurons, such as PPL1-α′3, consistent with previous reports (Huang et al., 2024).

**Figure 3.**
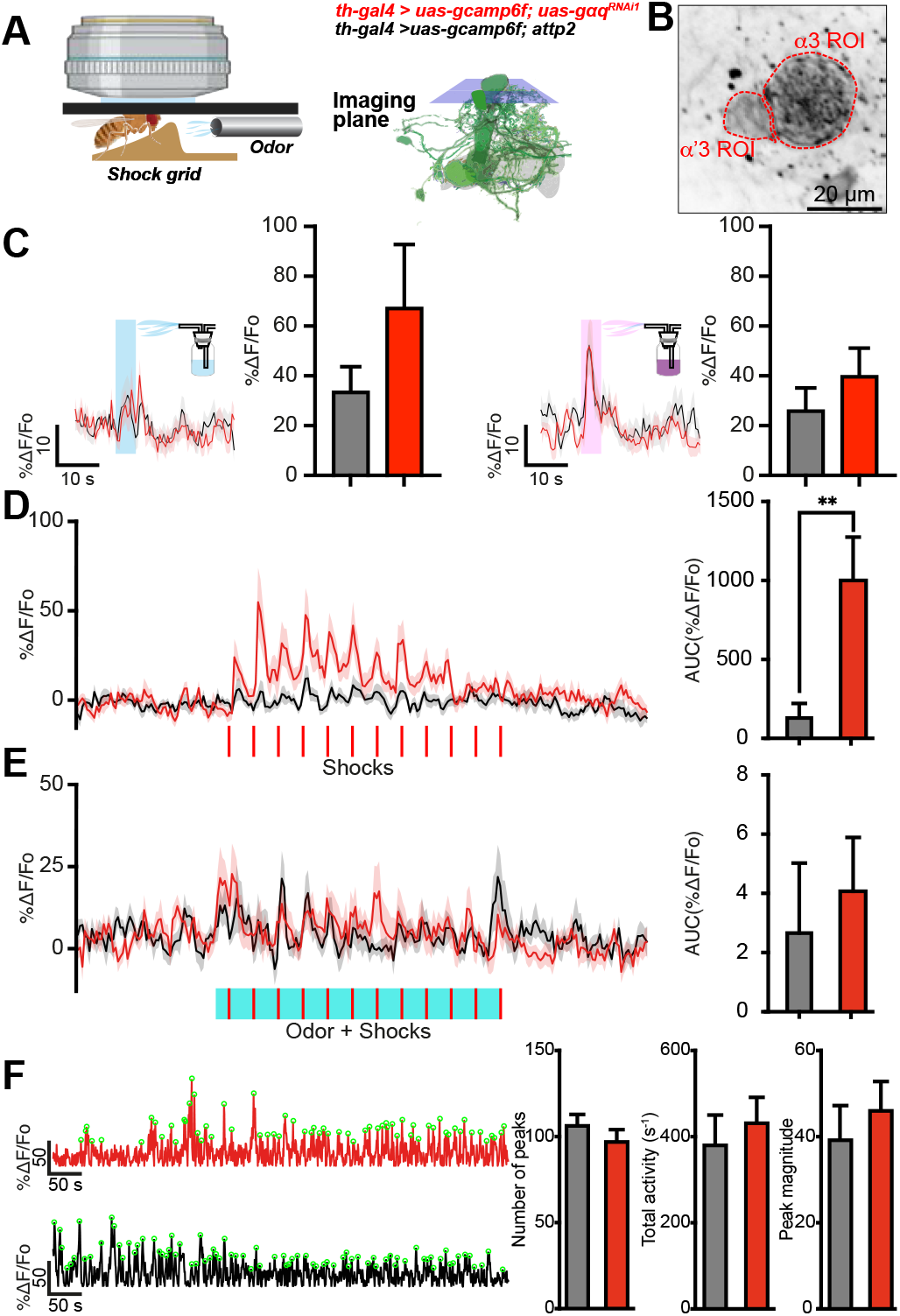
*gαq* knockdown in PPL1 DAN increases shock-evoked responses in PPL1-α3 terminals. (A) Left, schematic showing the in vivo functional imaging setup with a head-fixed fly resting on an electric shock grid and facing the odor delivery system. Middle, schematic of the cells and the imaging plane recorded. (B) Mean time series projection of baseline GCaMP6f driven by *th-gal4* in the PPL1-α3 and PPL1-α’3 projections. ROI is indicated by dotted red line. (C) GCaMP6f traces from PPL1-α3 terminals during odor presentation for 5 s, in control (black) and *gαq^RNAi^* (red) flies. Stimuli onset illustrated in the color shades. 4-MCH (blue) and 3-OCT (magenta) were used as odorants. Quantification of AUC shows no significant difference between groups. Mann–Whitney test; *n = 12-14*. Data are shown as mean ± SEM. (D) Shock-evoked GCaMP6f responses in PPL1-α3 terminals during 12 electric shocks (90 V, 1.25 s) in *gαq^RNAi^* (red trace) and control flies (black trace). Shock onsets are represented by the 12 red ticks below the traces. Traces show mean responses and shaded areas show the SEM. Right, trace quantifications show that *gαq* knockdown increases Ca^2+^ shock-evoked responses. Mann-Whitney test. *\*\*p = 0.0033. n = 14-16*. Data are shown as mean ± SEM. (E) GCaMP6f traces from PPL1-α3 terminals during an aversive conditioning protocol, with 60 s continuous odor presentation paired with 12 electric shocks in control (black) and *gαq^RNAi^* (red) flies. Stimuli onset illustrated below. Right, quantification shows no significant difference between groups. Mann–Whitney test. *p = 0.5267. n = 12-14*. Data are shown as mean ± SEM. (F) Knocking down *gαq* in *th-gal4* neurons does not alter ongoing activity in the PPL1-α3 terminals. Left. Representative traces of flies expressing the *gαq^RNAi^* transgene (red) and the control (black) in a ten-minute recording. Right. Ongoing activity is not statistically different in *gαq^RNAi^*when measuring number of peaks, total activity s-1 or peak magnitude. Two-tailed Mann-Whitney test. Data are shown as mean ± SEM.

We next examined whether *gαq* knockdown alters the neural response to the unconditioned stimulus (US) used during aversive conditioning. We recorded responses to 12 electric shocks (90 V, 1.25 s each), matching the stimulation protocol used for behavioral conditioning. *gαq* knockdown significantly increased shock-evoked Ca²⁺ transients in PPL1-α3 terminals compared with genetic controls (Fig. 3D). In contrast, when flies received the complete aversive conditioning protocol consisting of continuous odor presentation paired with 12 electric shocks, no significant difference in Ca²⁺ responses between genotypes was observed (Fig. 3E). We also examined ongoing PPL1-α3 activity over a 10-min period in the absence of external stimuli. No significant differences were detected between genotypes in any measure of ongoing activity, including peak number, total activity per second, or mean peak magnitude (Fig. 3F).

As an additional control, we simultaneously recorded activity from PPL1-α′3 terminals, which can be imaged in the same focal plane as PPL1-α3. This population provides an internal anatomical control because selective *gαq* knockdown in PPL1-α′3 did not alter memory performance in our behavioral experiments. Consistent with this behavioral result, *gαq* knockdown did not significantly alter any of the functional imaging parameters measured in PPL1-α′3 terminals (data not shown).

Together, these results indicate that *gαq* knockdown selectively enhances shock-evoked Ca²⁺ responses in PPL1-α3 terminals without altering odor-evoked responses, spontaneous activity, or responses in the behaviorally unaffected PPL1-α′3 population. These findings are consistent with a specific role for Gαq signaling in regulating the strength of the aversive US response in PPL1-α3 DANs.

## Identification of GPCRs mediating the Gαq-dependent regulation of memory

The experiments described above establish that Gαq signaling in PPL1 dopaminergic neurons regulates aversive memory and that this effect is associated with enhanced activity of specific PPL1-DAN compartments during aversive reinforcement. However, Gαq is a broadly utilized intracellular signaling pathway that can be activated downstream of numerous G protein-coupled receptors (GPCRs). Thus, the behavioral and physiological phenotypes produced by *gαq* knockdown do not, by themselves, identify the upstream receptor or receptors responsible for regulating memory through this pathway. Identifying the relevant GPCRs would provide a more specific entry point into the circuit and signaling mechanisms through which dopaminergic neurons regulate memory suppression.

To identify candidate GPCRs, we took advantage of previously published RNA-sequencing datasets generated from individual PPL1-DAN populations (Aso et al., 2019). We reasoned that GPCRs expressed at high levels in PPL1 neurons would represent strong candidates for upstream regulation of Gαq signaling in these cells. We therefore used the available transcriptomic datasets to identify GPCRs that are robustly expressed in PPL1-DANs (Aso et al., 2019), with particular emphasis on receptors expressed in the PPL1 populations implicated by our behavioral experiments. We then systematically tested candidate receptors using cell-type-specific RNA interference and asked whether knockdown of individual GPCRs reproduced the enhanced-memory phenotype observed following *gαq* knockdown. This approach allowed us to move from the broad manipulation of a common intracellular signaling pathway toward identification of the specific receptor pathways that may regulate memory through Gαq signaling in PPL1 DANs.

To identify GPCRs that may act upstream of Gαq to suppress aversive memory in PPL1 DANs, we analyzed previously published RNA-sequencing data from PPL1-DAN subpopulations (Aso et al., 2019). Based on detectable expression in PPL1 neurons, we selected 37 GPCRs as candidates for an initial RNAi screen. Candidate receptors were knocked down using the MB504B driver, which targets multiple PPL1 DANs (Aso et al., 2014), and aversive memory was assessed 3 h after training. Memory scores were normalized to the corresponding genetic controls (attP2 or attP40) for each RNAi line. This screen identified multiple GPCRs whose knockdown significantly enhanced memory performance. In contrast, knockdown of *fz2* and *op1R2* significantly reduced memory performance (Fig. 4A). Thus, the screen identified several GPCRs whose activity in PPL1 DANs may normally contribute to the suppression of aversive memory as Gαq does.

**Figure 4.**
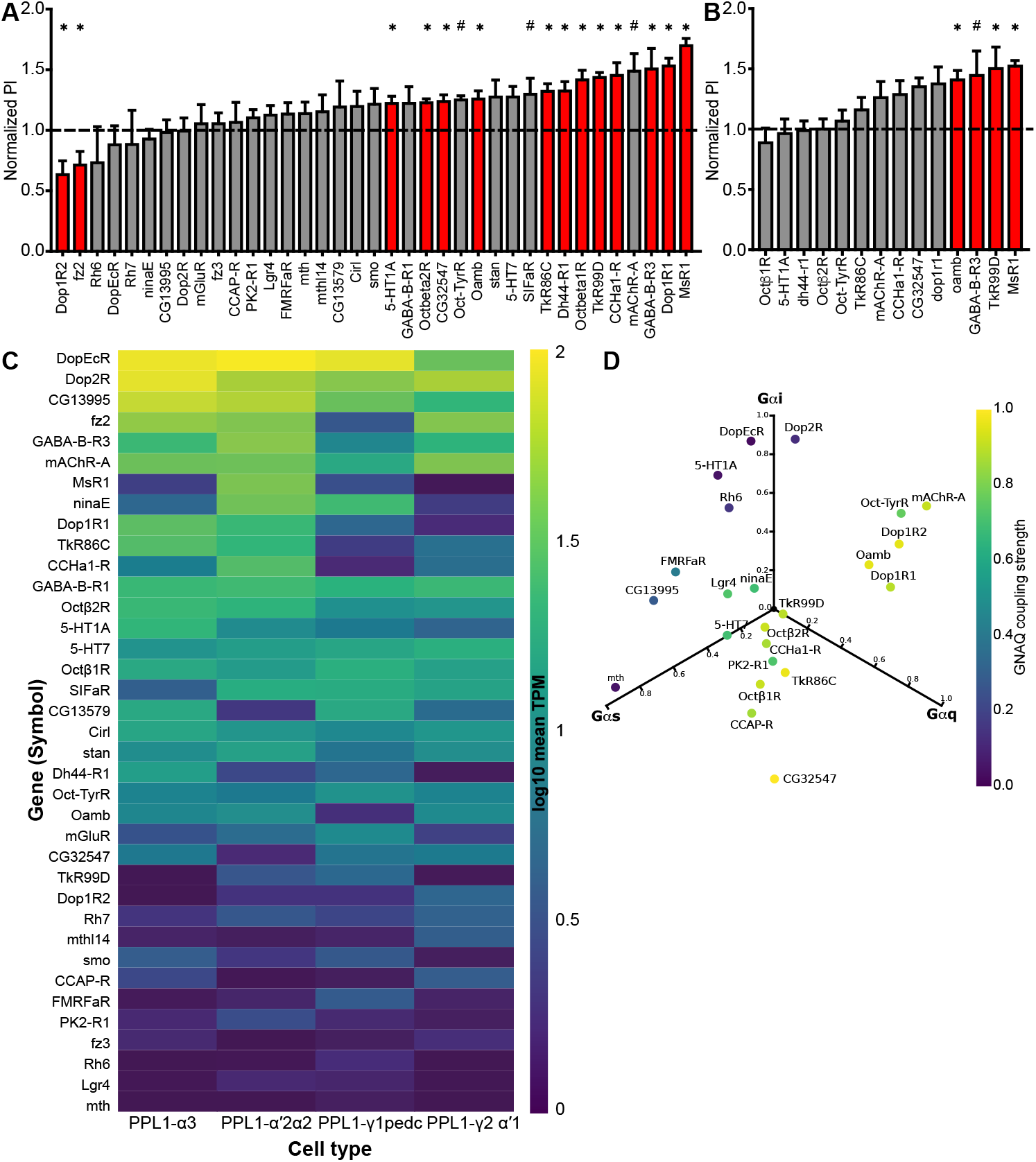
Identification of GPCRs as memory suppressors. (A) Normalized 3-h aversive memory scores for the GPCR knockdowns in PPL1 DANs using the MB504B split-*gal4* driver. Two-tailed Mann-Whitney tests comparing the receptor knockdown against its genetic control (attP2 or attP40). Normalized PI > 1 or < 1 indicates higher or lower memory than control, respectively. Asterisks indicate statistical significance. *\*p < 0.05*. (B) Normalized scores of the GPCR knockdowns in PPL1-α3 neurons after 3 h aversive memory. Receptors were screened based on yielding a significantly higher memory performance in the MB504B screen shown in A. (C) Heatmap of GPCR transcript expression across PPL1 dopaminergic neuron subtypes (RNA-seq data from Aso et al., 2019). Rows list GPCRs ranked by highest log mean TPM across PPL1 subtypes. Columns represent PPL1-α3, PPL1-α′2α2, PPL1-γ1pedc, and PPL1-γ2α′1. Color scale indicates log mean TPM. (D) G-protein coupling predictions (PRECOG) for the PPL1-expressed Class A GPCRs identified in (C). Each GPCR is plotted in a three-dimensional space defined by predicted coupling probabilities to Gαq (GNAQ), Gαs (GNAS), and Gαi (mean of GNAI1 and GNAI3). Marker color encodes Gαq coupling strength.

We next examined the expression of the candidate GPCRs across individual PPL1-DAN subtypes to determine whether the behavioral hits were expressed in the specific populations implicated by our *gαq* knockdown experiments. Transcript abundance (TPM) was compared across PPL1-α3, PPL1-α′2α2, PPL1-γ1pedc, and PPL1-γ2α′1 (Fig. 4C). Two dopamine receptors, *dop1R1* and *Dop1R2*, were among the most highly expressed GPCRs across these PPL1 populations, while the remaining candidates represented receptors for a diverse range of neuropeptides and neuromodulators. These data indicate that PPL1 DANs express a broad repertoire of GPCRs that could potentially regulate their function.

Because our experiments implicate Gαq signaling in the regulation of memory, we next asked whether the candidate receptors identified in the behavioral screen were predicted to couple to Gαq. We used PRECOG, a sequence-based predictor of GPCR–G protein coupling (Singh et al., 2019), to estimate the probability of Gαq coupling for the highly expressed candidate receptors. A three-dimensional scatter plot of predicted coupling to Gαq, Gαs, and Gαi revealed that several memory-enhancing hits, including mAChR-A and OAMB clustered with receptors predicted to have a high probability of Gαq coupling (Fig. 4D). This prediction is consistent with previous evidence that these receptors can signal through Gαq-dependent pathways and provided an additional criterion for prioritizing candidate receptors that could function upstream of Gαq (Himmelreich et al., 2017).

We next asked whether the GPCRs identified in the broad PPL1 screen also regulate memory specifically in PPL1-α3, the population in which *gαq* knockdown produced a robust memory phenotype and enhanced shock-evoked Ca²⁺ responses. PPL1-α3 has a distinct functional profile among PPL1-DANs, contributing relatively little to immediate aversive memory while supporting longer-lasting memory following spaced training, and is recruited during conditioning later than several early shock-responsive PPL1 subtypes (Aso and Rubin, 2016; Huang et al., 2024). Moreover, our expansion microscopy experiments revealed high levels of Gαq expression in this population. We therefore reasoned that GPCRs acting upstream of Gαq in PPL1-α3 should provide particularly strong candidates for mediating the enhanced-memory phenotype.

To test this, we re-screened 14 of the memory-enhancing GPCR candidates identified with MB504B using MB630B, which restricts RNAi expression to PPL1-α3 (Aso and Rubin, 2016). Aversive memory was again assessed 3 h after training. Knockdown of *msR1*, *tkR99D, oamb*, *cg32547, and dop1R1* significantly enhanced memory performance, whereas *gaba-b-R3* knockdown produced a trend toward enhanced memory (Fig. 4B). Thus, a subset of GPCRs identified in the broader PPL1 screen also suppresses aversive memory when their expression is selectively reduced in PPL1-α3, providing candidate upstream pathways through which Gαq signaling may regulate memory in this compartment.

## Functional characterization of GPCR knockdown effects on PPL1-α3 activity

We next asked whether knockdown of the candidate GPCRs altered PPL1-α3 terminal activity in a manner that could account for the enhanced-memory phenotype and whether any of the GPCR knockdowns reproduced the functional signature observed following *gαq* knockdown. We therefore quantified GCaMP6f signals in PPL1-α3 terminals during four experimental conditions: odor presentation, spontaneous activity, electric shock, and the complete odor–shock conditioning protocol. To facilitate comparison across genotypes and across different response measures, effect sizes were calculated as Hedges’ g relative to the corresponding matched genetic control.

We first examined odor-evoked activity in response to 4-MCH and 3-OCT. Odor responses were quantified as the area under the curve (AUC) during each 5-s odor presentation (Fig. 5A). Knockdown of *SIFaR* and *oamb* produced the largest changes in odor-evoked responses, whereas *gαq^RNAi^* produced non-significant changes in responses to both odors (Fig. 5A). These results indicate that several GPCRs identified in the behavioral screen can substantially alter sensory responses in PPL1-α3, although their effects are not necessarily similar to those produced by *gαq* knockdown.

**Figure 5.**
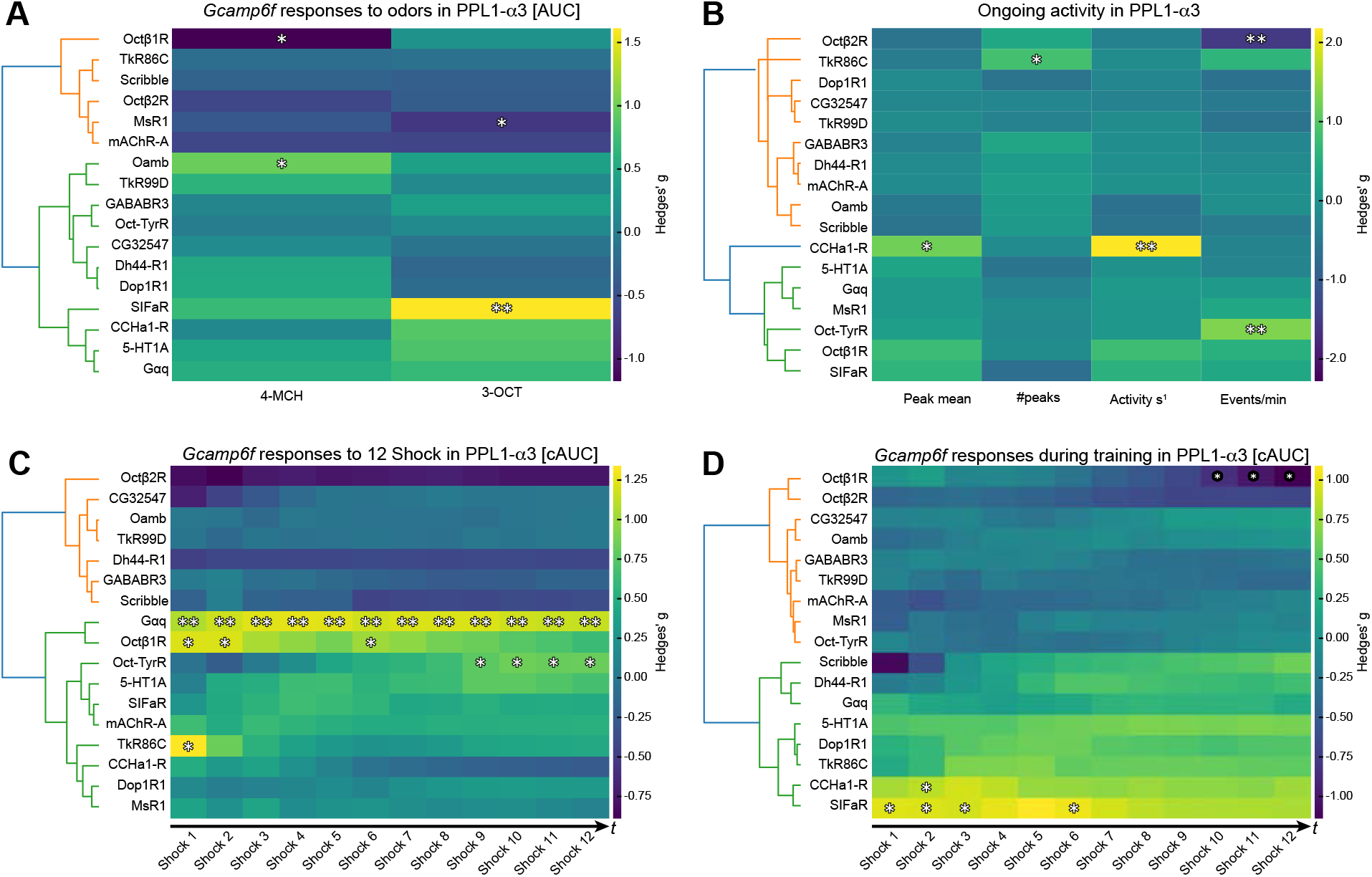
in vivo calcium imaging reveals gene-specific response profiles at PPL1-α3 terminals. (A) Heatmap of odor-evoked GCaMP6f responses (AUC) at PPL1-α3 terminals across GPCR, *scribble*, and *gαq* knockdown lines. Odors (4-MCH and 3-OCT) were delivered as 5 s pulses. Color scale indicates Hedges’ g. Asterisks indicate uncorrected two-tailed Mann-Whitney U significance. *\*p < 0.05, **p < 0.01*. (B) Ongoing activity at PPL1-α3 terminals. Columns show mean peak amplitude, number of peaks (prominence-filtered), activity per second (integrated ΔF/F per second), and events per minute (threshold crossings above mean + 2.5 SD). ΔF/F computed using a block-minimum baseline (10 s blocks). Color scale indicates Hedges’ g. Asterisks indicate uncorrected two-tailed Mann-Whitney U significance. *\*p < 0.05, **p < 0.01*. (C) Shock-evoked GCaMP6f responses (AUC) at PPL1-α3 terminals during 12 electric shocks (90 V, 1.25 s). Each column represents one shock event. Each trial was baselined to its own pre-shock window. Color scale indicates Hedges’ g. Asterisks indicate uncorrected two-tailed Mann-Whitney U significance. *\*p < 0.05, **p < 0.01*. (D) GCaMP6f responses during aversive conditioning (continuous 60 s odor paired with 12 electric shocks). Columns represent odor-anchored time bins aligned to each shock delivery within the odor presentation. Color scale indicates Hedges’ g. Asterisks indicate uncorrected two-tailed Mann-Whitney U significance. *\*p < 0.05, **p < 0.01*.

We next examined spontaneous PPL1-α3 activity in the absence of external stimulation. Ongoing activity was quantified using several measures, including mean peak amplitude, number of peaks, activity per second, and events per minute (Fig. 5B; see Methods). Among the GPCR knockdowns, *CCHa1-R* produced the most pronounced increase in baseline activity, with an effect size of Hedges’ g> 2 for activity per second. In contrast, *gαq^RNAi^* produced relatively modest changes across measures of spontaneous activity (Fig. 5B).

We then examined responses to the aversive unconditioned stimulus by presenting 12 electric shocks using the same stimulation protocol employed during behavioral conditioning. Shock-evoked activity was quantified as AUC over the complete 5-s response window for each shock and as cumulative AUC across all 12 shocks (Fig. 5C). The temporal profiles of shock responses differed substantially across knockdowns. *gαq^RNAi^* produced a consistent enhancement of shock-evoked responses across trials, accompanied by a modest adaptation-like decline. In contrast, *oct-tyrR* knockdown produced a progressive increase in responses across shocks, consistent with sensitization, whereas *ccha1-R* knockdown produced a progressive decline in response magnitude (Fig. 5C). These distinct response profiles indicate that the candidate GPCRs do not uniformly regulate PPL1-α3 responses to aversive stimulation.

Notably, *oamb^RNAi^* did not substantially alter shock-evoked Ca²⁺ responses in PPL1-α3, despite producing a robust behavioral memory phenotype. This dissociation suggests that OAMB-dependent suppression of memory may occur through mechanisms that do not involve modulation of the acute US-evoked Ca²⁺ response in PPL1-α3 terminals. Thus, the behavioral effects of GPCR knockdown cannot be explained solely by changes in the magnitude of shock-evoked activity.

Finally, we examined PPL1-α3 responses during the complete conditioning protocol, in which continuous odor presentation was paired with the 12-shock sequence (Fig. 5D). Interestingly, the enhanced shock response observed following *gαq* knockdown was not evident during conditioning. The mean effect size for *gαq^RNAi^* during the odor–shock protocol was close to zero (Hedges’ *g* = −0.01), in contrast to the strong enhancement observed when shocks were presented in isolation (Fig. 5C). This difference suggests that the effect of Gαq knockdown on PPL1-α3 activity is context dependent and is particularly evident in response to the aversive US in the absence of concurrent odor stimulation. Alternatively, the enhanced responses to shock during conditioning might be masked by the Ca²⁺ transients produced by the odor response.

## Testing Gαq-dependent octopamine signaling in dopaminergic neurons

Although none of the GPCR knockdowns fully reproduced the combination of behavioral and functional imaging phenotypes observed following *gαq* knockdown, several GPCRs produced significant effects on aversive memory. This lack of complete phenocopy is not unexpected, given that Gαq is a common intracellular signaling pathway that can integrate signals from multiple GPCRs. Disrupting *gαq* therefore has the potential to affect signaling downstream of several receptors simultaneously, whereas knockdown of an individual GPCR would be expected to perturb only one component of this broader signaling network. We therefore focused on the behavioral effects of individual GPCRs as a means of identifying signaling pathways that may contribute to Gαq-dependent regulation of memory.

One of the strongest candidates identified in our screen was *oamb*, which encodes a receptor for the neuromodulator octopamine. Although *oamb* knockdown produced an enhanced-memory phenotype, its effects on PPL1-α3 Ca²⁺ activity did not closely resemble those produced by *gαq* knockdown. Nevertheless, the behavioral phenotype raised the possibility that octopaminergic signaling could act through Gαq in dopaminergic neurons. We therefore sought to test more directly whether octopamine can engage Gαq-dependent intracellular Ca²⁺ signaling in DANs.

To monitor Ca²⁺ release from intracellular stores, we expressed the ER-targeted calcium indicator ER-GCaMP (de Juan-Sanz et al., 2017; Handler et al., 2019) in dopaminergic neurons and first validated the ability of the reporter to detect Gαq-dependent Ca²⁺ release in cell bodies. As a positive control, we bath-applied caffeine, which can open ryanodine receptors and promote Ca²⁺ release from intracellular stores. Caffeine application produced a robust response in ER-GCaMP fluorescence in dopaminergic neuron cell bodies, confirming that the reporter was responsive to pharmacologically induced Ca²⁺ release from the ER (Fig. 6A–B).

**Figure 6.**
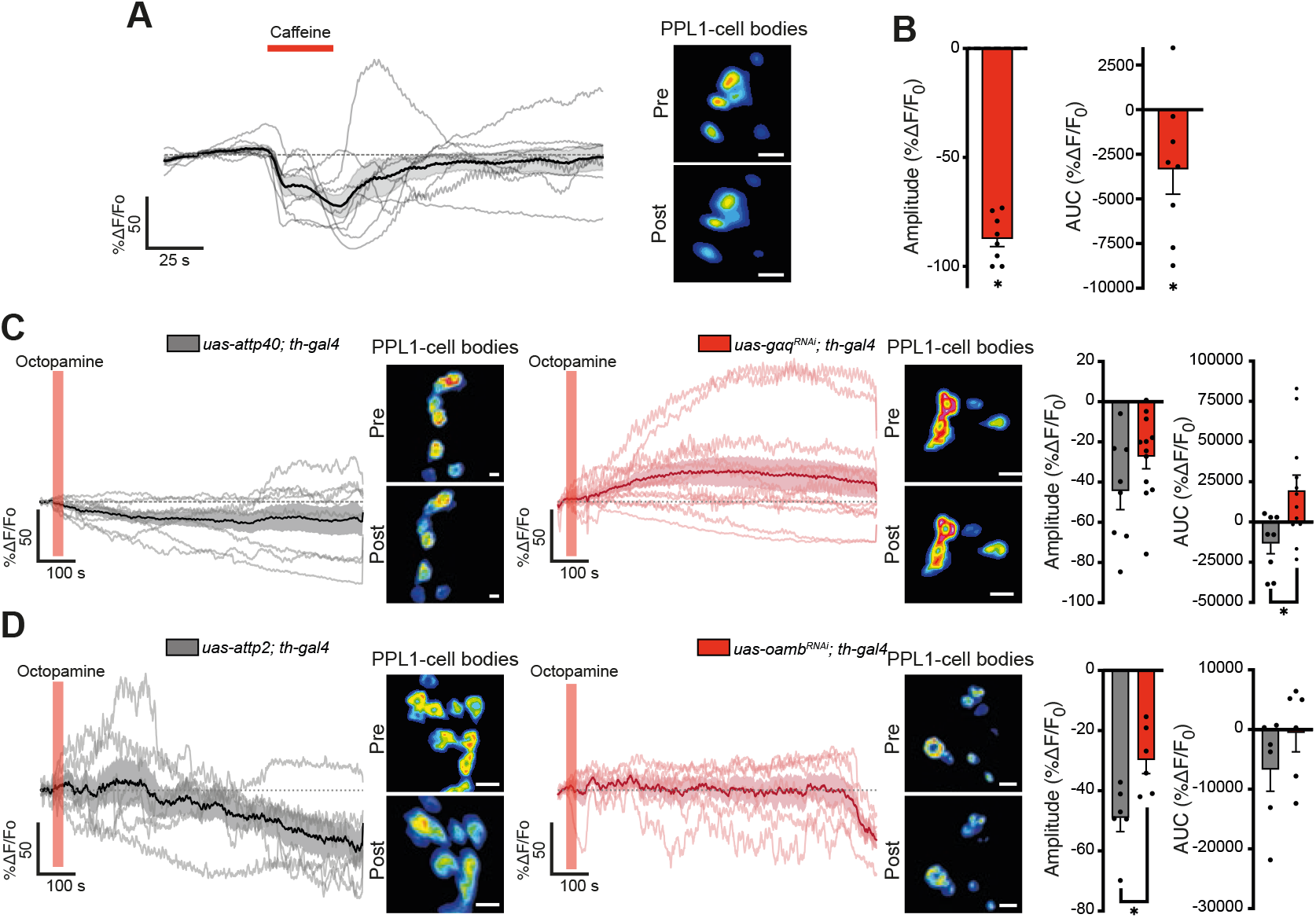
*gαq* and *oamb* knockdown impair octopamine-induced ER Ca²⁺ release in dopaminergic neuron cell bodies. (A) ER-GCaMP6-210 traces from PPL1 DAN cell bodies in *ex vivo* whole-brain preparations perfused with 10 mM caffeine (red bar). Individual flies shown as thin gray lines. Black trace with shaded area shows group mean ± SEM. *n = 8 flies*. Application of caffeine evoked robust ER Ca²⁺ response (mean amplitude = −87.1% ΔF/Fo), serving as a positive control for ER store mobilization in PPL1 DANs. Scale bar: 10 µm. (B) Amplitude and AUC quantification for caffeine. Each dot represents one fly. Representative ER-GCaMP6-210 images show baseline (top, 0–30 s average) and post-caffeine (bottom, 60 s–end average) intensity projections. Two-tailed paired t-test, pre vs post caffeine. Amplitude: *\*p < 0.0001*. AUC: *\*p = 0.049*. Scale bar: 10 µm. (C) Octopamine perfusion (1 mM, red bar) in *gαq* knockdown and control flies. Left: control traces. *n = 8, black*. Right: *gαq^RNAi^* traces. *n = 12, red*. *gαq* knockdown impairs sustained ER Ca²⁺ depletion, yielding a significantly less negative AUC compared to controls. Two-tailed Mann-Whitney U. *\*p = 0.025*. No significant difference was detected in amplitude. *p = 0.181*. Scale bar: 10 µm. (D) Octopamine perfusion in *oamb* knockdown and control flies. Left: control traces. *n = 6, black*. Right: *oamb^RNAi^* traces. *n = 6, red*. *oamb* knockdown results in a lower amplitude. \**p = 0.015*. No significant difference was detected in AUC. *p = 0.240*. Data shown as mean ± SEM. Scale bar: 10 µm.

We next asked whether octopamine similarly alters ER Ca²⁺ signals and whether this response depends on Gαq and OAMB receptor signaling. Octopamine was bath-applied while ER-GCaMP fluorescence was recorded from dopaminergic neuron cell bodies in control flies and flies expressing *gαq^RNAi^* (Fig. 6C) or *oamb^RNAi^* (Fig. 6D). Octopamine produced a significant ER-GCaMP response in control neurons, and this response was significantly altered following both *gαq* and *oamb* knockdown (Fig. 6C, D). These results suggest that octopamine can engage intracellular Ca²⁺ signaling in dopaminergic neurons and that this response is sensitive to disruption of Gαq signaling.

This experiment provides a more direct functional test of the possibility that octopaminergic signaling engages Gαq through OAMB in dopaminergic neurons than our initial behavioral and terminal-imaging experiments. However, several caveats should be considered. First, bath application of octopamine does not reproduce the spatial or temporal dynamics of endogenous octopamine release and does not establish that OAMB is the only receptor mediating the observed response. Second, ER-GCaMP reports changes in intracellular Ca²⁺ associated with ER stores but does not directly measure Gαq activation or receptor–G protein coupling. Finally, the lack of correspondence between *oamb^RNAi^* and *gαq^RNAi^* in PPL1-α3 terminal imaging indicates that OAMB cannot, on the basis of these experiments alone, account for the full physiological phenotype produced by *gαq* knockdown. Nevertheless, the differential octopamine-evoked ER-Ca²⁺ response following *gαq* knockdown provides functional evidence that octopaminergic signaling in dopaminergic neurons can engage a Gαq-dependent intracellular Ca²⁺ pathway.

## Concluding remarks

The genetic basis of memory suppression in *Drosophila* has typically been studied in Kenyon cells, where dopamine acting through DAMB and Gαq limits memory expression. Our findings show that Gαq signaling also acts within the dopaminergic neurons that convey reinforcement during conditioning. Knocking down *gαq* in PPL1 dopaminergic neurons enhanced aversive memory from acquisition through 24 h, and this effect did not depend on developmental disruption. When examined using split-*gal4*, memory was enhanced when *gαq* was knocked down specifically in PPL1-α3, PPL1-α′2α2, and PPL1-γ1pedc compartments. These results indicate that Gαq-dependent suppression is not confined to the postsynaptic side of the mushroom body circuit but is also built into the dopaminergic reinforcement pathway itself. Gαq in these neurons constrains how strongly aversive experiences are encoded, as shown by enhanced acquisition and increased shock-evoked Ca²⁺ responses in PPL1-α3 terminals.

The behavioral consequences are also distinctive. *gαq* knockdown enhanced both anesthesia-sensitive and anesthesia-resistant memory without altering long-term memory consolidation. It selectively impaired reversal learning and the loss of an original memory after competing training, while retroactive and proactive interference remained largely intact. Gαq in dopaminergic neurons therefore tunes memory strength and behavioral flexibility rather than acting as a general forgetting mechanism. This graded, process-specific profile helps explain how a single suppressor can influence several stages of memory and may help reconcile why different suppressors produce divergent effects on memory and flexibility. Finally, our screens implicate multiple GPCRs, including receptors for octopamine, tachykinin, and acetylcholine, as modulators of memory in PPL1 neurons. Octopamine engages Gαq-dependent Ca²⁺ release in dopaminergic neurons, suggesting that this pathway integrates diverse neuromodulatory inputs.

Several caveats apply. Gαq is a shared effector for many GPCRs, so no single receptor knockdown reproduced the full behavioral and imaging phenotype. The dissociation between *oamb* knockdown and PPL1-α3 terminal activity shows that enhanced shock-evoked Ca²⁺ cannot be the sole mechanism. Moreover, the enhanced shock response was not evident during odor–shock pairing. Receptor coupling was inferred from prediction and ER Ca²⁺ imaging with bath-applied octopamine, which does not directly measure Gαq activation or reproduce endogenous release. Future work should test endogenous ligand release and the downstream effectors that link Gαq to dopaminergic output.

## Acknowledgments

We would like to acknowledge the MakerHub at Georgetown for support building custom made tools for this project. This work was supported by National Institute of Health grants R01GM147917 (IC-S); F31GM157963 (PS); and the Brain and Behavior Research Foundation for 30442.

## Author contributions

Conceptualization: IC-S; behavioral experiments and analysis: JCM-C; imaging experiments and analysis: PS; microscopy: JCM-C; funding acquisition: IC-S, and PS; supervision: IC-S; writing: JCM-C, IC-S.

## Materials and methods

### Drosophila husbandry

Flies were raised on standard cornmeal medium at room temperature. Unless otherwise noted, crosses and experimental flies were maintained at 25 °C, 70% relative humidity, and a 12 h light– dark cycle. Table S1, shows the different stock lines used in this study.

### Immunoprecipitation and Mass spectrometry

To identify additional protein components that might be involved in memory suppression, we used the scrb::GFP^CA07683^ (Buszczak et al., 2007) gene trap line as bait to pull down Scribble-associated proteins. An equivalent number of wild type heads (endogenous, non-tagged Scribble) were processed in parallel as controls. Approximately 1,000 adult flies were anesthetized, collected, and flash-frozen in liquid nitrogen. Frozen flies were vortexed to separate heads from bodies and passed through a series of sieves (U.S. Standard Sieve Series) to isolate heads. Collected heads were transferred to microcentrifuge tubes and homogenized in 200 µL of CHAPS buffer supplemented with protease inhibitors (cOmplete, Roche) using a motorized pestle, while maintained on ice. An additional 800 µL of CHAPS buffer was added to reach a final volume of 1 mL. Homogenates were centrifuged at maximum speed for 10 minutes at 4 °C, and the supernatant was collected and clarified by a second centrifugation step. Extracts were repeated three times, corresponding to a total of ∼3,000 fly heads. Four separate samples were prepared and processed separately. Fly head extracts were then pre-cleared by incubation with control beads pre-coupled with rabbit IgG (Cell Signaling Technology 2729S) for 4 h at 4 °C with gentle agitation. Beads were pelleted by centrifugation (3,000 ×g, 2 min, 4 °C), and the supernatant was collected. An aliquot (20 µL) was reserved as the input sample, mixed with 6× sample buffer, boiled for 10 min, and stored at −20 °C.

For immunoprecipitation, pre-cleared extracts were incubated overnight at 4 °C with anti-GFP agarose beads (Abcam, ab69314) under constant agitation. Beads were pelleted (3,000 ×g, 2 min, 4 °C), and a fraction of the supernatant was collected as the output sample. Beads were washed three times with CHAPS buffer and bound proteins were eluted using 0.1 M glycine-HCl (pH 2.0), followed by immediate neutralization with Tris (pH 8.5). Eluates were pooled, concentrated using centrifugal filters (3 kDa cutoff), and buffer-exchanged into CHAPS buffer. Final samples were prepared with 6× sample buffer, boiled, and used for downstream analyses.

Immunoprecipitated proteins were fractionated by SDS–polyacrylamide gel electrophoresis. The gel was Coomassie-stained for 1 h at room temperature with shaking, followed by de-staining in water overnight. The gel bands were treated with 25 mM DTT followed by 55 mM iodoacetamide, and subjected to trypsin digestion with ProteaseMax Surfactant Trypsin Enhancer for one hour at 50°C. The peptide pools from each sample were acidified and desalted through Zip-Tip µC18 tip columns. Prior to mass spectrometry analysis the samples were reconstituted in 5 µL of 0.1% formic acid and 5 µL were loaded into the system. Each sample was analyzed by an Orbitrap Fusion™ Tribrid™ Mass Spectrometer (Thermo Fisher Scientific) coupled to an EASY-nLC 1000 system. Protein identification was carried out using Sequest algorithm, allowing oxidation (Met) and deamination (Q) as variable modifications. Other settings included carbamidomethylation of Cys as fixed modification, three missed cleavages, and mass tolerance of 10 ppm and 0.02 Da for precursor and fragment ions, respectively. MS/MS raw files were searched against a *Drosophila* database.

### Olfactory conditioning

Behavioral experiments used 2–5-day-old flies. Aversive olfactory conditioning was performed with minor modifications of established protocols (Beck et al., 2000). Two groups of ∼60 flies were trained reciprocally in a standard training tube with the following sequence: 30 s air, 60 s CS+ odor paired with 12 electric shock pulses (90 V, 1.25 s each), 30 s air, 60 s CS− odor without shock, 30 s air. Odors were 4-methylcyclohexanol (MCH) and 3-octanol (OCT), diluted in mineral oil. Memory was tested in a T-maze by presenting CS+ and CS− simultaneously in opposite arms for 2 min. Scores were calculated as a performance index. For time points other than 3 min, flies were returned to food vials after training until testing.

For shock-number acquisition curves, flies were trained as described above except the CS+ included 1, 2, 3, 6, or 12 shock pulses delivered evenly across the 60 s CS+ period such that the final pulse coincided with the end of odor presentation. For voltage acquisition curves, 12 shocks were delivered during the CS+ at 10, 20, 45, or 90 V.

For reversal learning paradigm, flies were first trained with the standard protocol (CS+ = MCH, CS− = OCT), followed by 1 min of air, then trained with the reversed contingency (CS+ = OCT, CS− = MCH). Memory was tested immediately after reversal training. For reversal learning with third odor, flies were trained as described for reversal learning (CS+ = OCT, CS-= BEN; reciprocal CS+ = MCH, CS-= BEN) followed by 1 min of air and the reverse odor-shock contingency (CS+=BEN, CS-=OCT; reciprocal CS+ = BEN, CS-= MCH). The memory test was performed using the odor first used as CS+ and a third odor (i.e., OCT and MCH) not experienced during training.

For retroactive interference experiments, flies were trained using the standard conditioning protocol with OCT and MCH. Then flies were re-trained 1.5 h after the first training session with a second odor combination using benzaldehyde (BEN) and methylsalicylate (MES). The memory test was performed 3 h after initial training using OCT and MCH. For proactive interference experiments, flies were trained using the standard conditioning protocol with BEN and MES, exposed to 1 min of air, and then re-trained using OCT and MCH. The memory test was performed immediately after the second training session using OCT and MCH.

For cold shock experiments, the cold shock was administered 2 h after conditioning for 2 min. Flies were transferred to glass vials at approximately 0 °C, then returned to food vials at 25 °C for recovery. Memory was tested 3 h after conditioning.

For TARGET (McGuire et al., 2003) experiments, flies were raised at 18 °C. One-day-old adults were shifted to either 18 °C or 30 °C for 3 days before behavioral testing. For all conditions, flies were placed at 25 °C for 1 h prior to conditioning and testing.

For long-term memory experiments, flies received five rounds of conditioning with either 15 min rest intervals between sessions (spaced) or no rest intervals (massed). Memory was tested 24 h after training.

### Immunostaining

Adult brains were dissected and immunolabeled based on standard methods with modifications. Briefly, male and female flies were anesthetized on ice or with carbon dioxide gas. Flies were rinsed once in 70% ethanol, washed and dissected in ice-cold S2 medium (Schneider’s Insect Medium, Sigma). Immediately after dissection, each brain was transferred to a 1.7 mL Eppendorf LoBind microcentrifuge tube containing 1 mL of 1% paraformaldehyde in S2 medium on ice. Samples were rotated at a fixed speed (24 rpm) overnight at 4 °C. After 16-24 h, fixative was aspirated and replaced with 1 mL of ice-cold PAT3 (0.5% Triton X-100, 0.5% bovine serum albumin in 1X PBS), then mixed by hand for a few seconds. PAT3 was discarded and brains were washed twice more in PAT3, with each wash performed at room temperature for 1 h. For blocking, brains were incubated in 1 mL of 3% normal goat serum (NGS) in PAT3 for 1.5 h at room temperature. Brains were then incubated in 1 mL of primary antibody solution prepared in blocking buffer at room temperature for 3 h, then overnight at 4 °C. The brains were washed in PAT3 three times (as described above). Then brains were incubated with secondary antibodies for 3 h at room temperature, followed by 5 days at 4 °C. Brains were washed again three times in PAT3, once in 1X PBS, and mounted on slides between 102 μm spacers in VectaShield mounting medium. Images were collected with a Leica TCS SP8 II confocal fluorescence microscope using a 20X (Leica Microsystems, HC APO L 20x/1.00) upright water-immersion lens. Primary antibodies used include: mouse monoclonal anti-Gαq/11 (1:75, Santa Cruz Biotechnology, cat #sc-515689, RRID: AB_2940775), rabbit polyclonal anti-Dlg1 (1:50, Boster Bio, cat #DZ01287, RRID: N/A), rabbit polyclonal anti-Scribble (1:50, Cervantes-Sandoval et al, 2016, RRID:AB_2571725). Secondary antibodies used were donkey anti-rabbit IgG (H+L) highly cross-adsorbed conjugated to CF488A (1:400, Biotium, Cat# 20015, RRID: AB_10559669) and donkey anti-mouse IgG (H+L) conjugated to CF633 (1:400, Biotium, Cat# 20124-1, RRID: AB_10853607).

### Expansion Microscopy

The *Drosophila* expansion protocol and associated reagents were adapted from (Asano et al., 2018; Bullard et al., 2024). For antibody-labeled samples, immunostaining was performed as described above prior to expansion. Acryloyl-X, SE (AcX; Invitrogen cat. no. A20770) was prepared by mixing AcX with dimethyl sulfoxide (1:100) to make the anchoring solution, which was previously stored at –20 °C with a desiccant. Immediately before use, the AcX/DMSO stock was diluted 1:100 in 1X PBS and the brains were incubated in this solution overnight at room temperature (>6 h) without shaking, then washed twice in cold 1X PBS (15 min per wash) in the dark. Samples were kept on ice thereafter until the gelling solution was prepared.

Gelling stocks (Stock X, 4-Hydroxy-TEMPO (4HT), ammonium persulfate (APS), and N,N,N′,N′-tetramethylethylenediamine (TEMED)) were thawed on ice. APS stock was prepared less than a week before use because reactivity decreases with long-term storage. Components were combined in a pre-chilled 1.7 mL Eppendorf Lo-Bind tube in the following order and ratio: Stock X, 4HT, TEMED, and APS (47:1:1:1). The gelling solution was vortexed briefly and returned to ice to minimize premature polymerization.

After replacing the previous buffer, the gelling solution was added to samples (500 μL per 6 to 10 brains) and incubated at 4 °C for 30 min in the dark. To limit agitation during transfers, gelation chamber slides were kept on a cold, flat, transferable surface. A 200 μL pipette tip was trimmed by removing ∼3 mm from the narrow end to transfer Drosophila brains onto the chamber. Brains were oriented to lie flat without wrinkles and embedded in gelling solution by adding 20–30 μL around the tissue. A coverslip was placed on the spacers and additional gelling solution was added until the chamber was filled without bubbles. Polymerization was carried out at 37 °C for about 3 h. After polymerization, gels were trimmed with a scalpel or razor blade around each brain to generate individual gel samples.

Digestion buffer aliquots stored at –20 °C without Proteinase K (ProK) were thawed immediately before use. Digestion buffer (2970 μL) was added to 35 mm polystyrene Petri dishes, and ProK (800 U/mL) was added (30 μL) to achieve a final dilution of 1:100. Samples were digested at room temperature in the dark for 14–16 h. Digestion was stopped by replacing digestion buffer with 1X PBS. Using two clean coverslips or scalpels, gels were transferred from 1X PBS to a dry Petri dish. Double-distilled water was then added and gels were incubated at room temperature for 20 min in the dark. This wash was repeated two additional times, after which the gels were larger and clearer. Post-expansion samples were imaged with the upright microscope mentioned above.

### In vivo functional calcium imaging

In vivo functional imaging was performed as previously described in animals without anesthesia (Shah and Cervantes-Sandoval, 2025). For in vivo Ca^2+^ imaging, GCaMP6f was co-expressed with RNAi lines under *th-gal4*. Head-fixed flies were imaged on a Leica TCS SP8 II confocal microscope using a 20x water-immersion objective. Time-series recordings were acquired at 2 Hz from PPL1-α3 synaptic terminals. Stimulus protocols included ongoing activity (10 min, no stimuli), shock-only (12 shocks at 90 V, 1.25 s each, 5 s ISI), odor-only (5 s pulses of 4-MCH and 3-OCT), and aversive conditioning (60 s continuous odor paired with 12 shocks). Baseline ΔF/Fo was computed from the first 27.5 s.

Ongoing activity metrics were computed from ΔF/Fo traces normalized to a block-minimum baseline (10 s blocks). Peaks were detected based on their height relative to surrounding values (minimum prominence = 0.25 × trace range), and mean peak amplitude was the average ΔF/Fo across detected peaks. Number of peaks was the count of events meeting this criterion. Activity per second was the integrated ΔF/Fo divided by recording duration. Events per minute was the rate of upward threshold crossings (mean + 2.5 SD).

Calcium imaging data were analyzed using custom scripts written in Python and MATLAB. Fluorescence time series were extracted from the PPL1-α3 or PPL1-α’3 ROI and converted to ΔF/Fo (%). For shock-only and odor-shock paradigms, Fo was computed from the first 55 timepoints (0–27.5 s at 2 Hz), which precede odor onset (30 s) and shock onset (33.75 s); this baseline was applied to the entire trace. For odor-only paradigms, Fo was computed separately for each odor epoch using the 10 s immediately preceding odor onset (odor 1: 20–30 s; odor 2: 55–65 s). Responses were quantified using two analysis windows per shock: a cycle window covering the full 5 s training unit (shock onset to next onset), and a shock window covering only the 1.25 s shock delivery period. For odor-shock conditioning, cycle windows were defined as odor-anchored 5 s bins spanning the 60 s odor epoch. For odor-only recordings, responses were quantified over each 5 s odor pulse.

Within each analysis window, response magnitude and dynamics were quantified using five metrics computed on ΔF/Fo (%): AUC (trapezoidal integral over the window), peak (maximum ΔF/Fo), latency to peak (time from window onset to peak), decay half-time (time from peak to 50% recovery), and plateau ratio (mean ΔF/Fo in the second half divided by the first half of the window). Cumulative AUC was computed as a running sum of per-shock AUC across the 12 shocks. AUC slope was computed by linear regression of per-shock AUC against shock number (1–12).

Each fly was treated as one independent observation. Group differences were summarized as Hedges’ g effect sizes, computed as the bias-corrected standardized mean difference between experimental and control groups. Positive g indicates larger responses in the experimental group. Hypothesis testing used two-sided Mann-Whitney U tests. Hedges’ g and p-values were compiled into matrices with rows corresponding to genotypes and columns to analysis windows and visualized as heatmaps. Genotypes were hierarchically clustered using Euclidean distance on the vector of Hedges’ g values across columns, with Ward linkage.

### Ex vivo ER-GCaMP imaging

Endoplasmic reticulum Ca^2+^ dynamics were assayed ex vivo using ER-GCaMP6-210 (Handler et al., 2019) expressed in DANs under *th-gal4*. Adult brains were dissected in physiological saline and maintained under continuous perfusion (2 mL/min). Time-series recordings were acquired at 2 Hz. Octopamine (1 mM) was applied by switching the inlet solution after a 30 s baseline period. Caffeine (10 mM) was used as a positive control for ER Ca²⁺ release. To account for drift, a mean intensity projection of the time-series stack was generated, and somatic regions of interest (ROIs) were drawn on this averaged image around individually resolved PPL1 dopaminergic neuron cell bodies. Each brain was treated as one biological replicate. ex vivo preparations were used because ER-GCaMP6-210 signal was detectable in PPL1 DAN cell bodies but not in axonal terminals. ΔF/Fo was calculated using the mean fluorescence of the 0–30 s pre-drug baseline period. Metrics were computed over the post-drug analysis window (60 s to end of recording, corresponding to 30 s after drug onset to allow for perfusion delivery). Per-fly traces were computed as the median ΔF/Fo across all ROIs. Sample sizes: *gαq*, n = 8 control, 12 experimental; *oamb*, n = 6 control, 6 experimental. For display, traces were smoothed with a 10-frame moving average. All statistics were computed on unsmoothed data.

### RNA-seq expression analysis

GPCR expression in PPL1 subtypes was analyzed using cell type-specific transcriptomic data from (Aso et al., 2019). Transcript-level expression values (TPM) were extracted from the supplementary dataset for four PPL1 cell types: PPL1-α3, PPL1-α’2α2, PPL1-γ1pedc, and PPL1-γ2α’1. A set of 37 GPCRs corresponding to the receptors screened behaviorally was selected from the full dataset. For genes with multiple transcript isoforms, expression was computed as the mean TPM across isoforms within each cell type. GPCRs were ranked by mean expression across PPL1 subtypes, and values were log-transformed (log10(TPM + 1)) for heatmap visualization.

### G protein coupling predictions

Predicted G protein coupling profiles were obtained using PRECOG (Singh et al., 2019) (precog.russelllab.org), which predicts coupling probabilities of Class A GPCRs to individual human Gα subunits based on receptor sequence features. *Drosophila* GPCR protein sequences were identified via FlyBase and mapped to UniProt accession numbers. Of the 37 GPCRs in the behavioral screen, 24 were classified as Class A and returned valid coupling predictions for all three axes used in visualization. Nine receptors (including GABA-B-R1, GABA-B-R3, mGluR, and members of the Frizzled and adhesion GPCR families) did not return valid predictions, consistent with their classification outside Class A. PRECOG returns coupling probabilities for 11 human Gα subunits (GNAI1, GNAI3, GNAZ, GNAO1, GNA12, GNA13, GNAQ, GNA14, GNA15, GNAS, GNAL). Results were visualized in a three-dimensional scatter plot with axes representing predicted coupling to Gαq (GNAQ), Gαs (GNAS), and Gαi (mean of GNAI1 and GNAI3), with marker color encoding GNAQ coupling strength.

---

**Figure S1.**
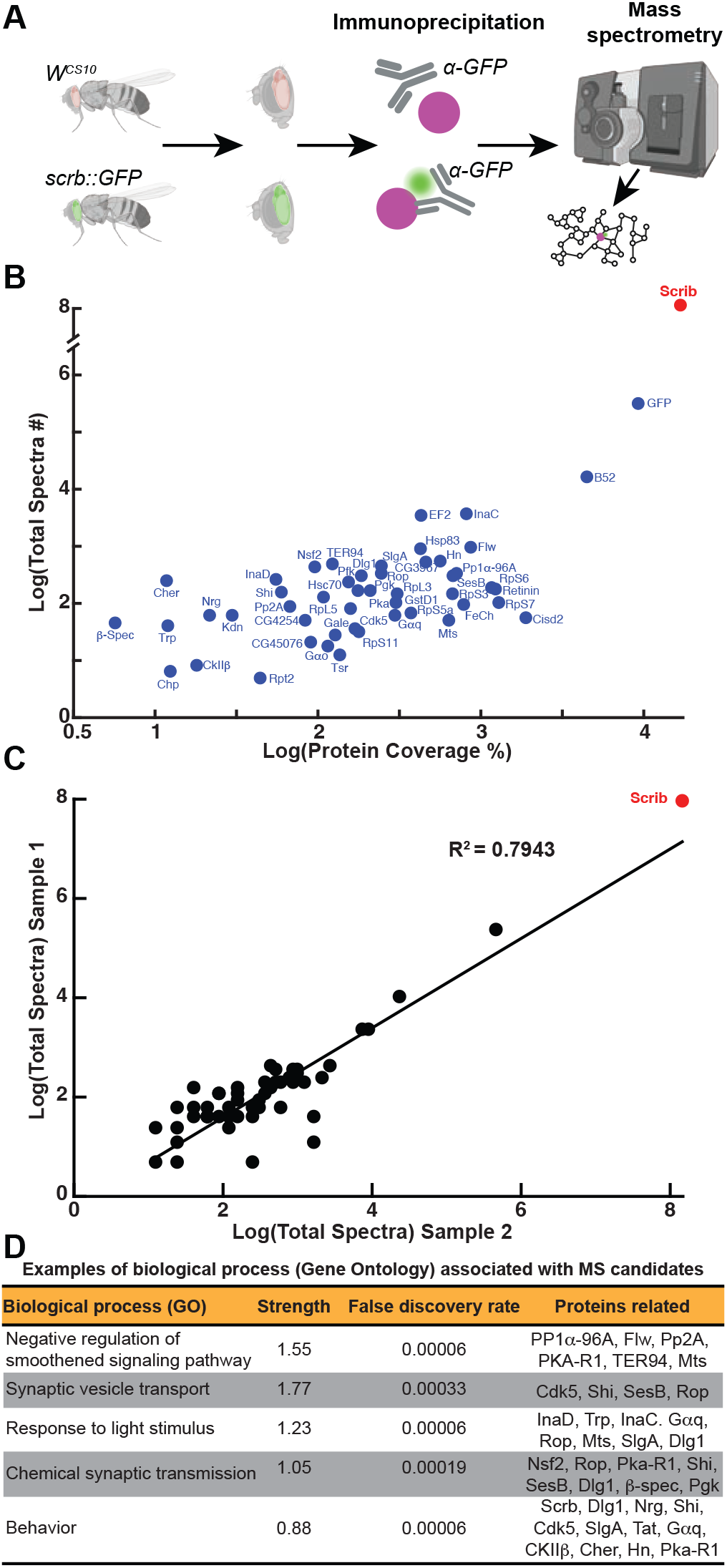
Co-IP/MS to discover proteins interacting with Scribble. (A) Diagram of experimental design. (B) Dot plot showing the average log (Total Spectra) and log (Protein Coverage) for the 54 MS candidates (including Scribble in red). (C) Correlation between candidates detected in two out of four different MS samples. (D) Five relevant biological processes (gene ontology) associated with the 54 MS candidates predicted by the STRING database. “Strength” denotes the ratio between the number of proteins in the network that are annotated with a term and the number of proteins that the database expected to be annotated with this term in a random network of the same size.

**Figure S2.**
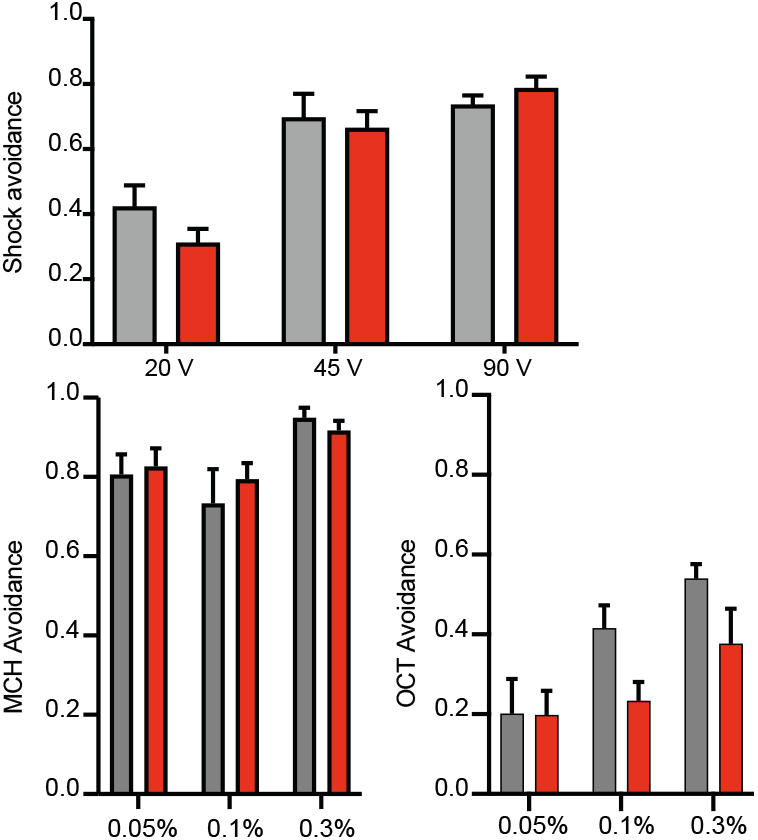
Sensory acuity and shock reactivity are normal in *gαq^RNAi^* flies. (A) Shock avoidance at 20, 45, and 90 V in *th-gal4*>*gαq^RNAi^*and genetic controls. No significant differences were observed at any voltage. Two-tailed Mann-Whitney test. 20 V: *p = 0.138, n = 9– 11*; 45 V: *p = 0.402, n = 5*; 90 V: *p = 0.442, n = 7*. Data are shown as mean ± SEM. (B) Naive avoidance to 4-methylcyclohexanol (MCH) at 0.05%, 0.1%, and 0.3% concentrations in *th-gal4* > *gαq^RNAi^* and genetic controls. No significant differences were observed. Two-tailed Mann-Whitney tests. 0.05%: *p = 0.711, n = 7–9*; 0.1%: *p = 0.965, n = 8–10*; 0.3%: *p = 0.365, n = 8*. Data are shown as mean ± SEM. (C) Naive avoidance to 3-octanol (OCT) at 0.05%, 0.1%, and 0.3% concentrations in *th-gal4* > *gαq^RNAi^* and genetic controls. No significant differences were observed. Two-tailed Mann-Whitney tests; 0.05%: *p = 0.628, n = 6–7*; 0.1%: *p = 0.052, n = 5–6*; 0.3%: *p = 0.097, n = 7*. Data are shown as mean ± SEM.

**Table S1.** Experimental models: *Drosophila* strains.

| **Fly line name** | **Source** | **Identifier** |
| --- | --- | --- |
| P{w[+mC]=ple-GAL4.F}3 [th-gal4] | Bloomington *Drosophila* Stock Center | BDSC:8848 |
| P{GawB}elav[C155] | Bloomington *Drosophila* Stock Center | BDSC:458 |
| w*;;P{nSyb-GAL4.S} | Gift from Julie Simpson |  |
| P{UAS-Dicer2}; | Vienna *Drosophila* Resource Center | VDRC:60009 |
| P{tubP-GAL80ts}; |  | NA |
| P{y[+t7.7] w[+mC]=R52H03-p65.AD}attP40;<br>P{y[+t7.7] w[+mC]=ple-GAL4.DBD}attP2 [splitGal4-MB504B] | Bloomington *Drosophila* Stock Center | BDSC:68329 |
| P{y[+t7.7] w[+mC]=VT026773-p65.AD}attP40;<br>P{y[+t7.7] w[+mC]=R72B05-GAL4.DBD}attP2 [splitGal4 MB630B] | Bloomington *Drosophila* Stock Center | BDSC:68334 |
| P{y[+t7.7] w[+mC]=ple-p65.AD}attP40; P{y[+t7.7] w[+mC]=R72B05-GAL4.DBD}attP2 [splitGal4 MB065B] | Bloomington *Drosophila* Stock Center | BDSC:68281 |
| P{y[+t7.7] w[+mC]=R82C10-p65.AD}attP40/CyO;<br>P{y[+t7.7] w[+mC]=R26F01-GAL4.DBD}attP2 [splitGal4 MB304B] | Bloomington *Drosophila* Stock Center | BDSC:68367 |
| P{y[+t7.7] w[+mC]=R15B01-p65.AD}attP40;<br>P{y[+t7.7] w[+mC]=R26F01-GAL4.DBD}attP2 [splitGal4 MB296B] | Bloomington *Drosophila* Stock Center | BDSC:68308 |
| P{y[+t7.7] w[+mC]=R22B12-GAL4.DBD}attP2 PBac{y[+mDint2] w[+mC]=ple-p65.AD} VK00027 [splitGal4 MB320C] | Bloomington *Drosophila* Stock Center | BDSC:68253 |
| P{y[+t7.7] w[+mC]=R82C10-p65.AD}attP40;<br>P{y[+t7.7] w[+mC]=R50B03-GAL4.DBD}attP2 [splitGal4 MB058B] | Bloomington *Drosophila* Stock Center | BDSC:68278 |
| P{y[+t7.7]=CaryP}attP40 | Bloomington *Drosophila* Stock Center | BDSC:36304 |
| P{y[+t7.7]=CaryP}attP2 | Bloomington *Drosophila* Stock Center | BDSC:36303 |
| P{y[+t7.7] v[+t1.8]=TRiP.JF02765}attP2 [Nsf2 RNAi] | Bloomington *Drosophila* Stock Center | BDSC:27685 |
| P{y[+t7.7] v[+t1.8]=TRiP.JF03205}attP2 [Tat RNAi] | Bloomington *Drosophila* Stock Center | BDSC:28777 |
| P{y[+t7.7] v[+t1.8]=TRiP.HM05140}attP2 [Rop RNAi] | Bloomington *Drosophila* Stock Center | BDSC:28929 |
| P{y[+t7.7] v[+t1.8]=TRiP.JF02261}attP2 [Hn RNAi] | Bloomington *Drosophila* Stock Center | BDSC:29540 |
| P{y[+t7.7] v[+t1.8]=TRiP.JF01217}attP2 [chp RNAi] | Bloomington *Drosophila* Stock Center | BDSC:31276 |
| P{y[+t7.7] v[+t1.8]=TRiP.JF03402}attP2 [ter94 RNAi] | Bloomington *Drosophila* Stock Center | BDSC:31968 |
| P{y[+t7.7] v[+t1.8]=TRiP.HMS00796}attP2 [Hsp83 RNAi] | Bloomington *Drosophila* Stock Center | BDSC:32996 |
| P{y[+t7.7] v[+t1.8]=TRiP.HMS00014}attP2 [dlg1 RNAi] | Bloomington *Drosophila* Stock Center | BDSC:33620 |
| P{y[+t7.7] v[+t1.8]=TRiP.HMS01088}attP2 [Cisd2 RNAi] | Bloomington *Drosophila* Stock Center | BDSC:33749 |
| P{y[+t7.7] v[+t1.8]=TRiP.HMS01324}attP2 [Pfk RNAi] | Bloomington *Drosophila* Stock Center | BDSC:34336 |
| P{y[+t7.7] v[+t1.8]=TRiP.HMS00888}attP2 [Hsc70-1 RNAi] | Bloomington *Drosophila* Stock Center | BDSC:34527 |
| P{y[+t7.7] v[+t1.8]=TRiP.HMS01129}attP2 [Gao RNAi] | Bloomington *Drosophila* Stock Center | BDSC:34653 |
| P{y[+t7.7] v[+t1.8]=TRiP.HMS00058}attP2 [CkIIβ RNAi] | Bloomington *Drosophila* Stock Center | BDSC:34939 |
| P{y[+t7.7] v[+t1.8]=TRiP.GL00189}attP2 [Cdk5 RNAi] | Bloomington *Drosophila* Stock Center | BDSC:35287 |
| P{y[+t7.7] v[+t1.8]=TRiP.HMS01501}attP2 [cher RNAi] | Bloomington *Drosophila* Stock Center | BDSC:35755 |
| P{y[+t7.7] v[+t1.8]=TRiP.HMS01549}attP2 [sesB RNAi] | Bloomington *Drosophila* Stock Center | BDSC:36661 |
| P{y[+t7.7] v[+t1.8]=TRiP.HMS01631}attP40 [kdn RNAi] | Bloomington *Drosophila* Stock Center | BDSC:36740 |
| P{y[+t7.7] v[+t1.8]=TRiP.GL01039}attP2 [GstD1 RNAi] | Bloomington *Drosophila* Stock Center | BDSC:36818 |
| P{y[+t7.7] v[+t1.8]=TRiP.HMS00154}attP2 [shi RNAi] | Bloomington *Drosophila* Stock Center | BDSC:36921 |
| P{y[+t7.7] v[+t1.8]=TRiP.HMS01638}attP40 [Nrg RNAi] | Bloomington *Drosophila* Stock Center | BDSC:37496 |
| P{y[+t7.7] v[+t1.8]=TRiP.HMS01803}attP2 [flw RNAi] | Bloomington *Drosophila* Stock Center | BDSC:38336 |
| P{y[+t7.7] v[+t1.8]=TRiP.HMS01804}attP2 [mts RNAi] | Bloomington *Drosophila* Stock Center | BDSC:38337 |
| P{y[+t7.7] v[+t1.8]=TRiP.HMS02154}attP40 [Pp1α-96A RNAi] | Bloomington *Drosophila* Stock Center | BDSC:40906 |
| P{y[+t7.7] v[+t1.8]=TRiP.GL01174}attP2 [β-Spec RNAi] | Bloomington *Drosophila* Stock Center | BDSC:42802 |
| P{y[+t7.7] v[+t1.8]=TRiP.GL01514}attP2 [slgA RNAi] | Bloomington *Drosophila* Stock Center | BDSC:43172 |
| P{y[+t7.7] v[+t1.8]=TRiP.HMS01921}attP2 [PpaA-2B RNAi] | Bloomington *Drosophila* Stock Center | BDSC:43283 |
| P{y[+t7.7] v[+t1.8]=TRiP.HMC02429}attP2 [Gale RNAi] | Bloomington *Drosophila* Stock Center | BDSC:44496 |
| P{y[+t7.7] v[+t1.8]=TRiP.HMC03170}attP2 [inaD RNAi] | Bloomington *Drosophila* Stock Center | BDSC:52313 |
| P{y[+t7.7] v[+t1.8]=TRiP.HMC03646}attP40 [PkaR1 RNAi] | Bloomington *Drosophila* Stock Center | BDSC:52906 |
| P{y[+t7.7] v[+t1.8]=TRiP.HMC03757}attP40 [FeCH RNAi] | Bloomington *Drosophila* Stock Center | BDSC:55146 |
| P{y[+t7.7] v[+t1.8]=TRiP.HMC04693}attP40 [Retinin RNAi] | Bloomington *Drosophila* Stock Center | BDSC:57389 |
| P{y[+t7.7] v[+t1.8]=TRiP.HMJ22520}attP40 [fau RNAi] | Bloomington *Drosophila* Stock Center | BDSC:60351 |
| P{y[+t7.7] v[+t1.8]=TRiP.HMJ23244}attP40 [trp RNAi] | Bloomington *Drosophila* Stock Center | BDSC:61352 |
| P{y[+t7.7] v[+t1.8]=TRiP.JF01209}attP2 [Gaq RNAi] | Bloomington *Drosophila* Stock Center | BDSC:31268 |
| P{y[+t7.7] v[+t1.8]=TRiP.GL01048}attP2 [Gaq RNAi] | Bloomington *Drosophila* Stock Center | BDSC:36820 |
| P{y[+t7.7] v[+t1.8]=TRiP.HMJ30300}attP40/CyO [Gaq RNAi] | Bloomington *Drosophila* Stock Center | BDSC:63987 |
| P{y[+t7.7] v[+t1.8]=TRiP.HMS00534}attP2 [tsr RNAi] | Bloomington *Drosophila* Stock Center | BDSC:65055 |
| P{y[+t7.7] v[+t1.8]=TRiP.HMC06255}attP40 [CG42540 RNAi] | Bloomington *Drosophila* Stock Center | BDSC:65975 |
| P{y[+t7.7] v[+t1.8]=TRiP.JF01845}attP2 [CG13575 RNAi] | Bloomington *Drosophila* Stock Center | BDSC:25827 |
| P{y[+t7.7] v[+t1.8]=TRiP.HMS00299}attP2 [SIFaR RNAi] | Bloomington *Drosophila* Stock Center | BDSC:34947 |
| P{y[+t7.7] v[+t1.8]=TRiP.JF01852}attP2 [5-HT1A RNAi] | Bloomington *Drosophila* Stock Center | BDSC:25834 |
| P{y[+t7.7] v[+t1.8]=TRiP.JF01879}attP2 [FMRFaR RNAi] | Bloomington *Drosophila* Stock Center | BDSC:25858 |
| P{y[+t7.7] v[+t1.8]=TRiP.HMS00191}attP2 [mGluR RNAi] | Bloomington *Drosophila* Stock Center | BDSC:34872 |
| P{y[+t7.7] v[+t1.8]=TRiP.HMC06293}attP2 [Dop1R2 RNAi] | Bloomington *Drosophila* Stock Center | BDSC:65997 |
| P{y[+t7.7] v[+t1.8]=TRiP.JF02271}attP2 [GABA-B-R3 RNAi] | Bloomington *Drosophila* Stock Center | BDSC:26729 |
| P{y[+t7.7] v[+t1.8]=TRiP.JF02363}attP2 [Smo RNAi] | Bloomington *Drosophila* Stock Center | BDSC:27037 |
| P{y[+t7.7] v[+t1.8]=TRiP.JF02576}attP2 [5-HT7 RNAi] | Bloomington *Drosophila* Stock Center | BDSC:27273 |
| P{y[+t7.7] v[+t1.8]=TRiP.JF02967}attP2 [Oct-TyrR RNAi] | Bloomington *Drosophila* Stock Center | BDSC:28332 |
| P{y[+t7.7] v[+t1.8]=TRiP.JF02989}attP2 [GABA-B-R1 RNAi] | Bloomington *Drosophila* Stock Center | BDSC:28353 |
| P{y[+t7.7] v[+t1.8]=TRiP.JF03017}attP2 [CNMaR RNAi] | Bloomington *Drosophila* Stock Center | BDSC:28380 |
| P{y[+t7.7] v[+t1.8]=TRiP.HMJ21906}attP40 [CNMaR RNAi] | Bloomington *Drosophila* Stock Center | BDSC:57859 |
| P{y[+t7.7] v[+t1.8]=TRiP.JF03036}attP2 [CG32547 RNAi] | Bloomington *Drosophila* Stock Center | BDSC:28621 |
| P{y[+t7.7] v[+t1.8]=TRiP.JF03070}attP2 [Lgr4 RNAi] | Bloomington *Drosophila* Stock Center | BDSC:28655 |
| P{y[+t7.7] v[+t1.8]=TRiP.JF03208}attP2 [Dh44-R1 RNAi] | Bloomington *Drosophila* Stock Center | BDSC:28780 |
| P{y[+t7.7] v[+t1.8]=TRiP.JF03303}attP2 [PK2-R1 RNAi] | Bloomington *Drosophila* Stock Center | BDSC:29624 |
| P{y[+t7.7] v[+t1.8]=TRiP.JF01673}attP2 [PK2-R1 RNAi] | Bloomington *Drosophila* Stock Center | BDSC:31171 |
| P{y[+t7.7] v[+t1.8]=TRiP.JF01732}attP2 [Oamb RNAi] | Bloomington *Drosophila* Stock Center | BDSC:31233 |
| P{y[+t7.7] v[+t1.8]=TRiP.JF01338}attP2 [CCAP-R RNAi] | Bloomington *Drosophila* Stock Center | BDSC:31490 |
| P{y[+t7.7] v[+t1.8]=TRiP.HM04077}attP2 [Dop1R1 RNAi] | Bloomington *Drosophila* Stock Center | BDSC:31765 |
| P{y[+t7.7] v[+t1.8]=TRiP.HMC05200}attP40 [Dop1R1 RNAi] | Bloomington *Drosophila* Stock Center | BDSC:62193 |
| P{y[+t7.7] v[+t1.8]=TRiP.JF01438}attP2 [ninaE RNAi] | Bloomington *Drosophila* Stock Center | BDSC:31647 |
| P{y[+t7.7] v[+t1.8]=TRiP.JF02645}attP2 [Mth RNAi] | Bloomington *Drosophila* Stock Center | BDSC:27495 |
| P{y[+t7.7] v[+t1.8]=TRiP.HMC06074}attP40 [mthl13 RNAi] | Bloomington *Drosophila* Stock Center | BDSC:65203 |
| P{y[+t7.7] v[+t1.8]=TRiP.HMC06446}attP2 [mthl14 RNAi] | Bloomington *Drosophila* Stock Center | BDSC:67342 |
| P{y[+t7.7] v[+t1.8]=TRiP.HMC03348}attP2 [CG11318 RNAi] | Bloomington *Drosophila* Stock Center | BDSC:51792 |
| P{y[+t7.7] v[+t1.8]=TRiP.JF02275}attP2 [CG13995 RNAi] | Bloomington *Drosophila* Stock Center | BDSC:26733 |
| P{y[+t7.7] v[+t1.8]=TRiP.JF02663}attP2 [TkR99D RNAi] | Bloomington *Drosophila* Stock Center | BDSC:27513 |
| P{y[+t7.7] v[+t1.8]=TRiP.JF02680}attP2 [MsR1 RNAi] | Bloomington *Drosophila* Stock Center | BDSC:27529 |
| P{y[+t7.7] v[+t1.8]=TRiP.JF02748}attP2 [CCHa1-R RNAi] | Bloomington *Drosophila* Stock Center | BDSC:27669 |
| P{y[+t7.7] v[+t1.8]=TRiP.JF03059}attP2 [CG13579 RNAi] | Bloomington *Drosophila* Stock Center | BDSC:28644 |
| P{y[+t7.7] v[+t1.8]=TRiP.JF01572}attP2 [Octβ1R RNAi] | Bloomington *Drosophila* Stock Center | BDSC:31107 |
| P{y[+t7.7] v[+t1.8]=TRiP.JF01378}attP2 [fz2 RNAi] | Bloomington *Drosophila* Stock Center | BDSC:31390 |
| P{y[+t7.7] v[+t1.8]=TRiP.HM05266}attP2 [CG12290 RNAi] | Bloomington *Drosophila* Stock Center | BDSC:31873 |
| P{y[+t7.7] v[+t1.8]=TRiP.JF02160}attP2 [TkR86C RNAi] | Bloomington *Drosophila* Stock Center | BDSC:31884 |
| P{y[+t7.7] v[+t1.8]=TRiP.JF03415}attP2 [DopEcR RNAi] | Bloomington *Drosophila* Stock Center | BDSC:31981 |
| P{y[+t7.7] v[+t1.8]=TRiP.HMS01151}attP2 [Octβ2R RNAi] | Bloomington *Drosophila* Stock Center | BDSC:34673 |
| P{y[+t7.7] v[+t1.8]=TRiP.HMS00136}attP2 [Cirl RNAi] | Bloomington *Drosophila* Stock Center | BDSC:34821 |
| P{y[+t7.7] v[+t1.8]=TRiP.HMS01464}attP2 [stan RNAi] | Bloomington *Drosophila* Stock Center | BDSC:35050 |
| P{y[+t7.7] v[+t1.8]=TRiP.GL01057}attP2 [Dop2R RNAi] | Bloomington *Drosophila* Stock Center | BDSC:36824 |
| P{y[+t7.7] v[+t1.8]=TRiP.GLC01626}attP2 [fz3 RNAi] | Bloomington *Drosophila* Stock Center | BDSC:44468 |
| P{y[+t7.7] v[+t1.8]=TRiP.HMC02343}attP2 [mAChR-A RNAi] | Bloomington *Drosophila* Stock Center | BDSC:44469 |
| P{y[+t7.7] v[+t1.8]=TRiP.HMC04325}attP40 [Rh6 RNAi] | Bloomington *Drosophila* Stock Center | BDSC:56029 |
| P{y[+t7.7] v[+t1.8]=TRiP.HMC05183}attP40 [Rh7 RNAi] | Bloomington *Drosophila* Stock Center | BDSC:62176 |

